# Image-based screens identify regulators of endogenous Dvl2 biomolecular condensates

**DOI:** 10.1101/2025.01.11.632522

**Authors:** Antonia Schubert, Florian Heigwer, Christian Scheeder, Oksana Voloshanenko, Dominique Kranz, Franziska Ragaller, Nadine Winkler, Thilo Miersch, Barbara Schmitt, Melanie Kuhse, Daniel Gimenes, Diana Ordoñez-Rueda, Jennifer Schwarz, Frank Stein, Dirk Jäger, Ulrike Engel, Michael Boutros

## Abstract

Dishevelled (Dvl) proteins are essential transducers in Wnt signaling pathways, which have been implicated in development, stem cell maintenance, and human diseases such as cancer. Several studies have shown that Dvl proteins form dynamic biomolecular condensates. However, how cellular signals and cell states influence the formation of biomolecular condensates remains poorly understood. Here, we analyzed cells with endogenous Dvl2 condensates using image-based cell sorting in combination with phosphoproteomics and identified protein enrichment for Wnt/PCP signaling and the G2/M cell cycle transition. We then performed an image-based high-throughput screen to identify small molecule kinase inhibitors that affect Dvl2 liquid-liquid phase separation. Strikingly, CK1δ/ε inhibition blocked Dvl2 condensate formation. Its effect on Wnt signaling was modulated in genetic epistasis experiments with loss-of-function alleles of APC, Axin1, and MCC. Our study highlights the interplay between post-translational modifications and condensate dynamics, opening new avenues for research on their role in cellular signaling and disease intervention.

## INTRODUCTION

Cellular signal transduction relies on intracellular compartmentalization, as the localization of signaling factors ensures specific protein interactions and enables the spatiotemporal control of the intracellular signals. A spatial arrangement through compartmentalization, e.g., at the membrane or organelles, is critical for signaling specificity, timing, and the strength of responses (Su et al., 2021; Banani et al., 2017; Alberti et al., 2019). An example of a complex and tightly regulated pathway is Wnt signaling, which controls critical biological processes, such as embryogenesis, stem cell homeostasis, and tissue patterning (Clevers and Nusse, 2012; Zhan et al., 2017; Gammons and Bienz, 2018; Bugter et al., 2021).

Dishevelled (Dvl) proteins are central signaling hubs involved in β-catenin-dependent and -independent Wnt pathways (Boutros and Mlodzik, 1999; Wallingford and Habas, 2005; Paclíková et al., 2021; Gentzel and Schambony, 2017; Boutros et al., 1998; Mlodzik, 2016). The Dvl proteins have conserved DIX (*Dishevelled and Axin)*, PDZ (*Post synaptic density, Disc large, Zonula occludens-1)*, and DEP (*Dvl, Egl-10, Pleckstrin)* domains, as well as intrinsically disordered regions (IDRs) (Boutros and Mlodzik, 1999; Kang et al., 2022; Schaefer and Peifer, 2019). In β-catenin-dependent (“canonical”) Wnt signaling, Wnt ligands bind to a Frizzled (Fzd) receptor, recruiting LRP6 as co-receptor, which leads to Dvl phosphorylation by CK1ε (Cong et al., 2004; Bilic et al., 2007; Peters et al., 1999; Bryja et al., 2007). Dvl subsequentl yoligomerizes through self-interaction mediated by its DIX domain and interacts with self-associated Axin proteins, which leads to the inactivation of the cytosolic β-catenin destruction complex comprising Axin1, APC, CK1α, and GSK-3β, (Gammons et al., 2016; Kan et al., 2020; Schwarz-Romond et al., 2007). β-catenin moves into the nucleus, where it activates the transcription of Wnt effector genes *via* T-cell factor (TCF) and lymphoid enhancer factor (LEF) (Behrens et al., 1996). Other Wnt ligands, such as Wnt5a, have been described to signal independent of β-catenin, e.g., the Wnt/PCP and Wnt/Ca^2+^ pathways modulating cytoskeletal dynamics, cell polarity, cell adhesion and migration (Clevers and Nusse, 2012).

Several steps in Wnt pathways involve membrane-less subcellular structures (Schaefer and Peifer, 2019; Hartmann et al., 2024). Compartmentalization of signaling pathway components often occurs in membranous organelles such as the endoplasmic reticulum (ER), Golgi, and endosomes, or, as more recently described, in membrane-less subcellular structures now termed biomolecular condensates (Su et al., 2021; Banani et al., 2017). These membrane-less structures form through liquid-liquid phase separation (LLPS) driven by locally increased protein polymers involving intrinsically disordered regions (IDRs) or nucleic acids (Banani et al., 2017; Lyon et al., 2021). Local signals can dramatically change phase-separated systems (Banani et al., 2016; Brangwynne et al., 2015; Pak et al., 2016). The importance of biomolecular condensates has been demonstrated for many molecular processes, from signal transduction to gene regulation during development and in diseases such as cancer (Su et al., 2021; Boija et al., 2021; Sabari, 2020). However, the mechanisms that govern biomolecular condensates’ formation, regulation, and dissolution remain poorly understood. Mechanisms that influence the assembly and disassembly of biomolecular condensates are also essential to understand in the context of drug action mechanisms. Targets residing in biomolecular condensates may have different accessibility for drugs, and drugs that affect the formation of biomolecular condensates may have widespread unintended effects on cellular processes (Mullard, 2019; Chakravarty et al., 2022).

Previously, it has been shown that upon overexpression, Dvl can form Dvl-enriched “puncta” accumulations (Schwarz-Romond et al., 2005). These membrane-free clusters change dynamically and expand through fusion in a concentration-dependent manner (Schwarz-Romond et al., 2005; Smalley et al., 2005; Sear, 2007). Additionally, their role at physiological protein levels and their relevance for signaling transmission have been discussed in the past (Schwarz-Romond et al., 2005; Smalley et al., 2005). It was also proposed that the “signalosome” and the “destruction complex” represent biomolecular condensates (Schaefer and Peifer, 2019; Bienz, 2020; Li et al., 2020; Nong et al., 2021). Imaging-based approaches offer the possibility of visualizing biomolecular condensates in living cells and could be used to identify chemical or genetic perturbations that affect their formation. Applying novel methodologies, such as CRISPR-mediated genome engineering, optogenetic tools, and super-resolution imaging techniques, allowed us to demonstrate that components of Dvl, Axin, APC, and β-catenin containing membrane-less condensate in higher resolution and showed that Wnt signaling pathway components undergo phase separation *in vitro* and *in vivo* (Schubert et al., 2022; Nong et al., 2021; Kang et al., 2022; Lach et al., 2022a; Schaefer et al., 2018; Kunttas-Tatli et al., 2014). In Dvl proteins, IDR1 drives Dvl2 phase separation, with the DIX domain aiding in this process (Kang et al., 2022). We recently demonstrated at a single molecule resolution that endogenous Dvl2 can dynamically associate with supramolecular structures at the centrosome (Schubert et al., 2022). Dvl2 co-localized in these centrosomal condensates with other Wnt signaling mediators, including APC and Axin1. While these studies gave insights into the internal organization of these native Dvl2 condensates at single-molecule resolution and demonstrated their Wnt- and cell cycle-dependent formation, questions remained on their assembly regulation, signaling activity, composition, druggability, and physiological functions.

Here, we performed high-content image-based screening combined with image-enabled cell sorting and phosphoproteomic analyses to gain novel insights into biomolecular condensates and drug discovery. By examining changes of the phosphoproteome of condensate-positive cells, we observed increased Wnt/PCP pathway activity and disruptions in the G2/M phase transition. We assessed a near kinome-wide library of 628 small molecule kinase inhibitors to evaluate their effects on canonical Wnt signaling activation and Dvl2 condensate formation by means of a TCF4/Wnt luciferase reporter assay and an image-based readout, respectively. Notably, CK1δ/ε inhibition by PF670462 disrupted Wnt signaling and lowered Dvl2 condensate counts while affecting cell division. Phosphoproteomic analysis identified key phosphosites and Wnt scaffolds involved in Dvl2 condensation. Using CRISPR/Cas9-mediated knockouts of APC, AXIN1, and MCC, we elucidated their distinct roles in regulating Dvl2 condensation during active Wnt signaling.

## RESULTS

### Image-enabled sorting of cells based on the detection of endogenous Dvl2 condensates

Cell populations harbour a heterogeneous distribution of endogenous Dvl2 condensates, as shown by fluorescent imaging upon tagging the DVL2 gene with a mEos3.2 fluorophore (Schubert et al., 2022). Dvl2 dynamically co-localizes with other Wnt signaling components in supramolecular centrosomal protein assemblies. Because of the biophysical features of Dvl proteins and the presence of IDRs, these structures were proposed to represent biomolecular condensates (Fig. S1) (Gammons and Bienz, 2018; Schubert et al., 2022; Kang et al., 2022; Ma et al., 2020). Super-resolution imaging revealed the internal architecture and formation of these structures. However, it remains unclear how the dynamic formation of these structures is controlled and which cell states favor the formation of Dvl2 condensates (Schubert et al., 2022).

To better understand cell states that allow condensate formation, we intended to sort cells based on the presence or absence of endogenous Dvl2 condensates. To this end, we established an image-based method to detect condensates in live cells. We used image-enabled cell sorting (Schraivogel et al., 2022), combining flow cytometry with the detection of subcellular structures (Fig. 1A). In order to confirm the phenotype of the sorted fractions and evaluate the dynamics of protein condensates, cells were sorted into clear-bottomed 96-well plates (Fig. 1B, C). Imaging of cells collected directly into paraformaldehyde (PFA) at time 0 confirmed that all condensate-positive sorted cells showed condensates. In contrast, condensate-negatives sorted cells were devoid of them (Fig. 1C). We then performed time-course experiments with cells sorted into complete medium. Sorted cells in both fractions started to proliferate after a few hours, and we demonstrated that the condensate-negative cell pool began to form Dvl2 condensates, indicating that the propensity to form condensates is not fixed in a population. Interestingly, the proportion of condensate-positive cells in the sorted positive pool remained higher for at least 96 h, even as the cells proliferated. As previously reported, mitotic cells showed no condensates (Schubert et al., 2022). These results indicate that both condensate-positive and condensate-negative sorted cells represent cell states that can dynamically transition into one another, rather than being two fixed cell populations.

**Fig. 1:**
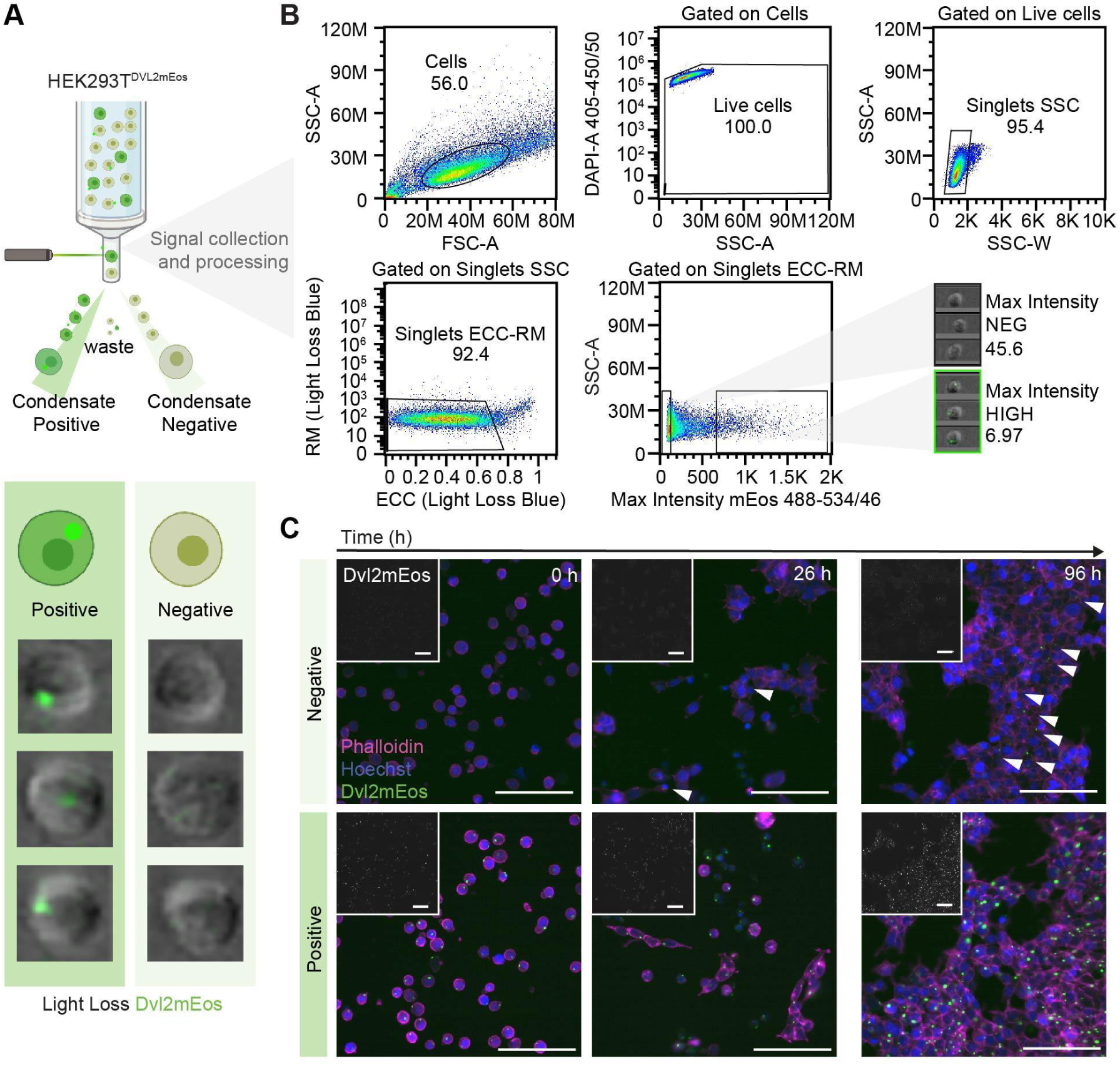
I**m**age**-enabled sorting of cells based on native Dvl2 condensates** (A) Schematic representation of the image-enabled cell sorting workflow. Dvl2 condensate-positive and -negative cells were sorted in 96-well clear-bottom wells for imaging and time-course experiments or into collection tubes for proteomic analyses. (B) Gating strategy for image-enabled cell sorting on Dvl2 condensates. RM: radial moment, ECC: eccentricity. (C) Time-course microscopy confirms the purity of the sorted fractions and demonstrates the dynamic formation of condensates in initially condensate-negative cells. After the indicated time points, the cells were fixed and labeled using Hoechst and DyLight Phalloidin 655. For T=0h, cells were sorted into PFA. Wide-field images were taken with an InCell Analyzer 6000 (20x magnification). Scale bar 100 µm.

### Cell-state analysis of Dvl2-condensate positive cells

In order to get insights into Dvl2-condensate positive and negative cells, we conducted quantitative proteomic profiling applying Tandem Mass Tag (TMT) multiplexing after image-based cell sorting. We split the lysate for each fraction, with one part subjected to phosphopeptide enrichment and another analyzed for global proteome changes. This approach allowed us to capture both phosphoproteome and full proteome dynamics across multiple cell sorting days (Fig. 2A). In total, we quantified 6270 proteins in the input and 9469 unique phosphopeptides in the condensate enriched fraction. Thirty-six proteins and 170 phosphopeptides were significantly regulated (fold-change ≥ 50 % and FDR ≤ 0.05 using limma, a moderated *t*-test) (Fig. 2B). Among the differentially regulated proteins, we detected a statistically significant increase in Dvl1 and Dvl2 proteins in the condensate-positive cell pools, while for other Wnt signaling scaffolds that have been shown to colocalize in centrosomal condensates, such as APC (Schubert et al., 2022) and CTNNB1 (Lach et al., 2022a), we did not detect changes in protein levels. Gene ontology enrichment analyses were performed using the clusterProfiler package in R to identify significantly enriched terms for cellular components, molecular function, and biological processes among proteins across different samples and conditions. Enriched terms (*p*.adjust < 0.05) were simplified to remove redundancy and visualized, with odds ratios indicating enrichment magnitude and -log_10_ adjusted *p*-values. The analyses revealed that in condensate positive cells, proteins associated with the Wnt pathway, such as Dvl1 and Dvl2, and proteins associated with microtubule-based processes and cytoskeleton organization, such as TBCD, ARL2, and CCNB1, were enriched. Interestingly, Wnt signaling-associated proteins were not statistically significantly differentially phosphorylated except CCDC88CC, also known as Daple. Instead, we detected differential phosphorylation of the proteins related to cell and nuclear division and cytoskeleton organization, including CEP131, CENPF, PRC1, BAIAP2, ANLN, WAC, and DDX3X (Fig. 2C). These results are consistent with our previous observation by superresolution microscopy that some condensates were observed near two mature CEP164-positive mother centrioles, a hallmark of cells preparing to enter mitosis (Schubert et al., 2022). These results suggest that Dvl2 condensate-positive cells represent a distinct cellular state characterized by G2/M phase progression. The enrichment of mitosis-related proteins and phosphorylation events, along with the localization of condensates at mature centrioles, supports the hypothesis that centrosomal Dvl2 condensates form during a cell cycle-dependent process.

**Fig. 2:**
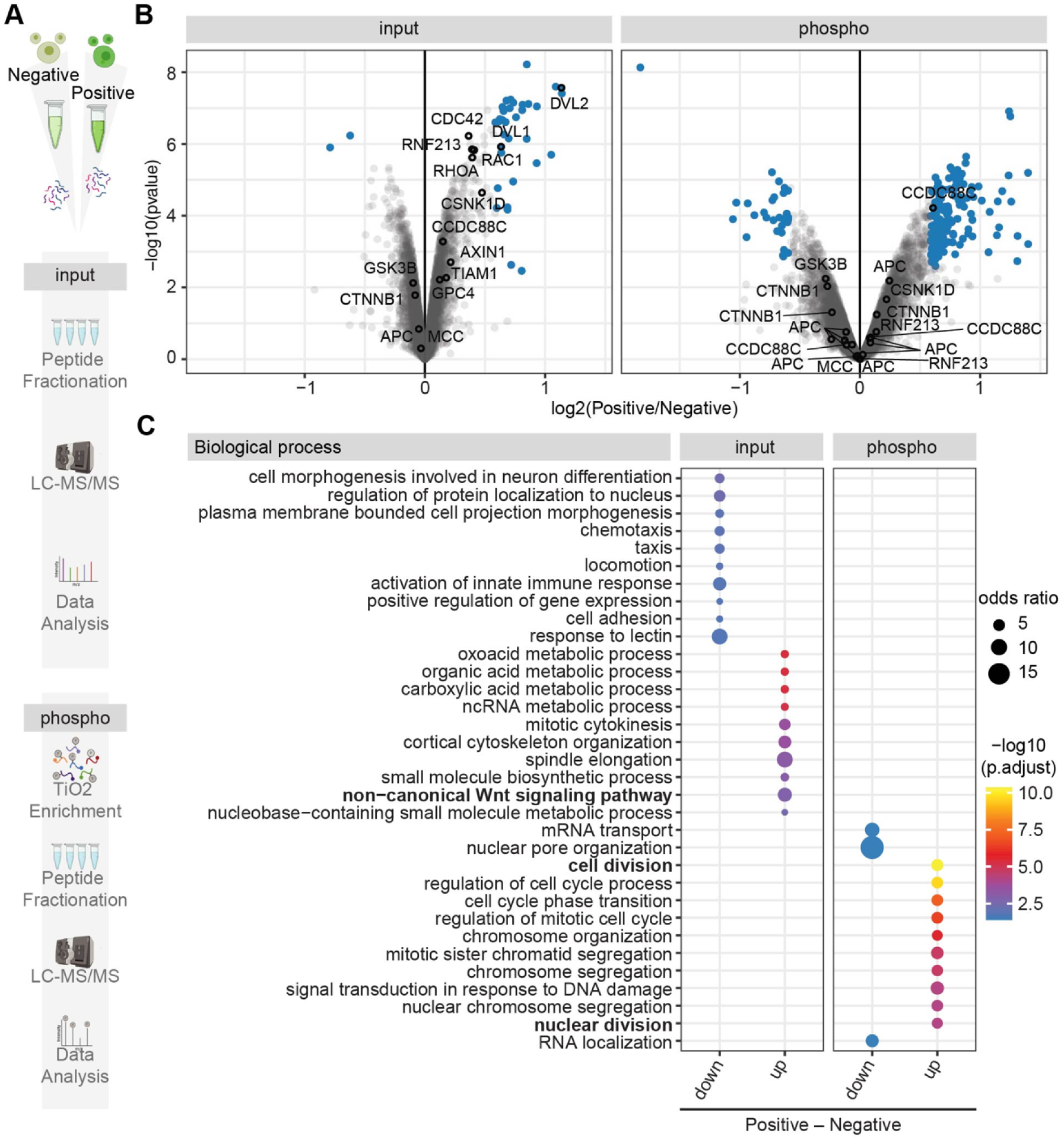
C**e**ll**-state analysis of Dvl2 condensate positive cells.** (A) Schematic presentation of the proteomic workflow. “Input” represents total protein, “phospho” the phosphorylated subset of proteins after enrichment. (B) Volcano plot of the identified proteins in the total (input) and phosphoproteomic (phospho) analyses. 6270 proteins were identified in the input and 9469 unique phosphopeptides in the condensate enriched fraction. 36 proteins and 170 phosphopeptides were significantly regulated (fold-change ≥ 50% and FDR ≤ 0.05 using limma, a moderated *t*-test). Top hits are highlighted in blue, and selected proteins are named. (C) GO enrichment analysis indicates an upregulation of proteins associated with the Wnt pathway and an upregulation of phosphoproteins associated with cell- and nuclear division in the Dvl2 condensate-positive *versus* -negative HEK293T^DVL2_mEos^ cells. The analysis was conducted using the clusterProfiler R package to identify significantly enriched biological processes. Terms with *p*.adjust < 0.05 were simplified to reduce redundancy and visualized as dot plots in ggplot2, with odds ratios (size of dots) representing enrichment magnitude and -log_10_ adjusted *p*-values (color) indicating significance.

### High-throughput image-based screening identifies modulators of Dvl2 condensates

Next, we performed an image-based near kinome-wide small molecule screen to investigate whether specific kinases can influence endogenous Dvl2 condensates. Post-translational modifications, such as phosphorylation, have recently been suggested to govern condensate formation and repression (Sridharan et al., 2022; Gerbich et al., 2020). Advances in image-based screening technologies have enabled phenotypic profiling for chemical and genetic perturbations. Microscopy-based profiling approaches focussing on single cell phenotypes have been used to explore the mechanisms of action, effectiveness, and toxicity of small molecules (Scheeder et al., 2018; Heigwer et al., 2023; Breinig et al., 2015; Piotrowski et al., 2017; Moshkov et al., 2023; Friese et al., 2019; Mattiazzi Usaj et al., 2020).

To investigate the role of phosphorylation in condensate biology and condensation of Wnt signaling components and to further understand the cellular state of condensate-positive and -negative cells, we performed an image-based kinase inhibitor screen (Fig. 3A). HEK293T^DVL2_mEos^ cells were seeded in 384-well plates and treated for 48 hours with a custom, kinase inhibitor library of 643 small molecules (15 spike-in controls and 628 samples) in two different concentrations (0.5 and 5µM). After 48 hours of treatment, cells were fixed, stained with Phalloidin and Hoechst, and imaged at 20x magnification using an automated microscope. Automated image analysis detected single cells based on DNA and F-actin staining, while Dvl2 condensates were identified by intensity-based thresholding of the FITC:mEos signal within segmented cell boundaries. The imaging screen was performed in three biological replicates, and a total of 66,528 images were analyzed.

**Fig. 3:**
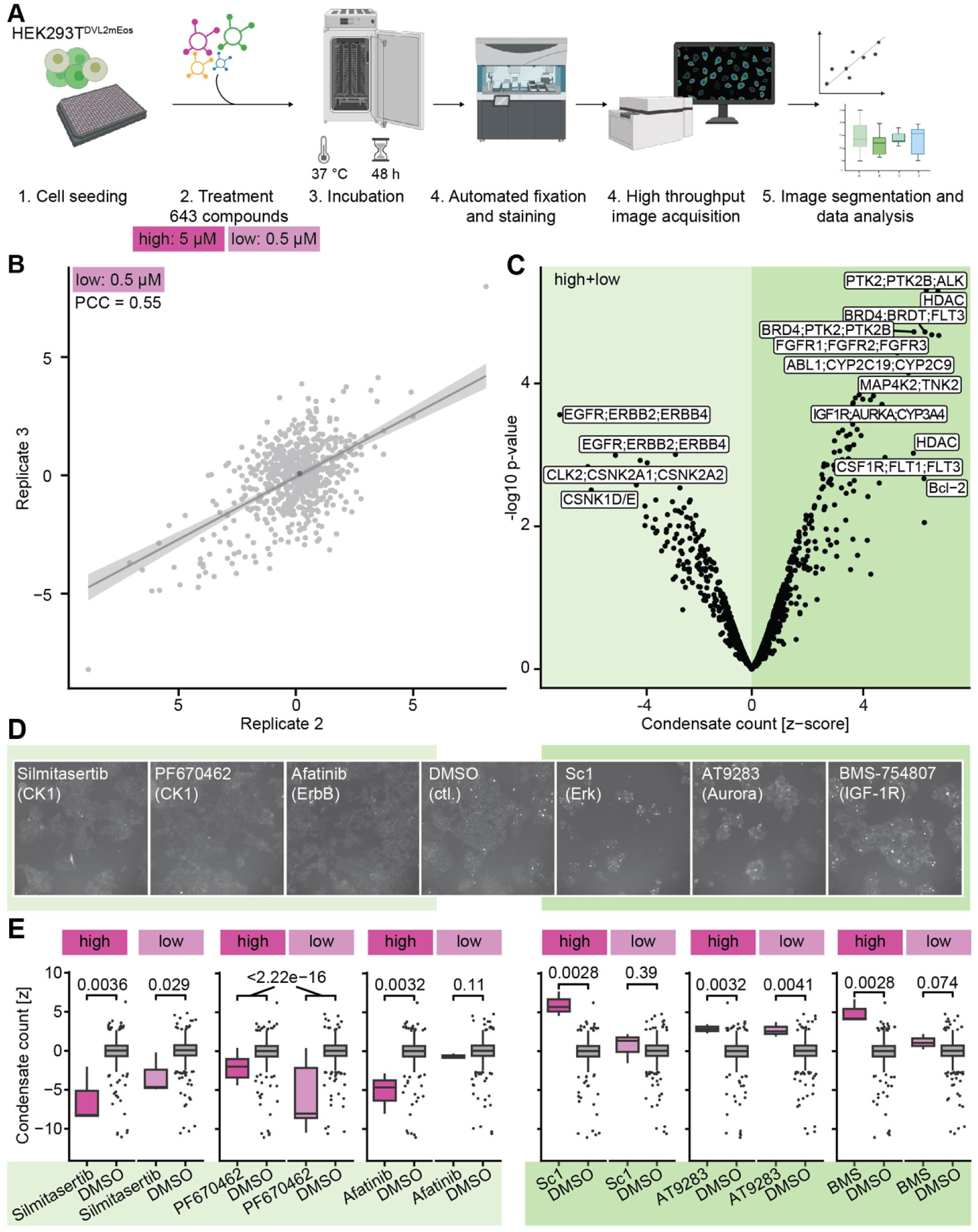
H**i**gh**-throughput image-based screening identifies inducers and inhibitors of Dvl2 condensate formation.** (A) Schematic presentation of the image-based small molecule screening workflow. HEK293T^DVL2_mEos^ cells were seeded into clear-bottom 384-well plates. After 24 h, the cells were treated with a custom small molecule inhibitor library at 0.5 μM (low) and 5 μM (high). After 48 h of treatment, the cells were fixed and labeled using with Hoechst and DyLight Phalloidin 655. Wide-field images were captured with an InCell Analyzer 2200 at 20x magnification, and automated image analysis was carried out as previously outlined (Schubert et al., 2022). N=3 biological replicates for each concentration. (B) Correlation of the screening results for relative condensate counts of the 2^nd^ and 3^rd^ replicate and cells treated with compounds at low concentrations demonstrates a high reproducibility of measurements (Pearson correlation coefficient [PCC] = 0.55). Condensates were counted per well and divided by the cell count of the respective well, the relative condensate count was then normalized per plate and z-score scaled. PCC values for additional replicates and concentrations are shown in Table S1. Light grey = compounds, dark grey = DMSO control. (C) Volcano plot showing inducers and inhibitors of Dvl2 condensates. The reduction of condensates is indicated by light green shading, while the induction of condensates is indicated by dark green shading. (D) Shown are representative images of inhibitors at 5 μM (high) concentrations reducing condensate counts (left of DMSO control) and inducing condensates (right of DMSO control). (E) Representative condensate counts of compounds shown in *(D)*. Boxplots display the median (black bar), the 25^th^ and 75^th^ percentile (box) ±1.5 times the interquartile range (whiskers). Outliers are plotted as separate dots outside the 25^th^ and 75^th^ percentile box. Statistical significance was estimated using the Wilcoxon signed rank test.

Activities of hit compounds were reproducible across three biological replicates (avg. PCC = 0.4, Fig. 3B and Table S1). Overall, 157 compounds modulated condensate formation at one or both concentrations tested (FDR<0.2, absolute effect size [z-score] >1). Among these, 55 compounds decreased and 101 increased averaged condensate counts at high, low, or both concentrations (Fig. 3C). Specifically, we found the JNK inhibitor Anisomycin, multiple PLK1 inhibitors (including NMS-P937, Ro3280, MLN0905), and inhibitors of casein kinases (Silmitasertib, SB239063, and PF670462 (see also Fig. 3D, E)) acting as inhibitors of condensate formation. Among the condensate count inducers, we identified GSK3 inhibitors (including LY2090314, CHIR-99021, SB415286), PTK2 inhibitors (PF00562271, PF573228, PF431396, PF562271), and multi-kinase inhibitors targeting MTOR/PIK3 (including LY3023414, GDC-0980, and GSK2126458). Strikingly, among the strong condensate inducers were multi-kinase inhibitors with known effects on cell cycle and cell proliferation as detected through cell counts, nuclear size, and nuclear eccentricity by our multiparametric feature analysis (Fig. S2). These included HDAC-, FLT3-, and Bcl2-inhibitors, which disrupt cell cycle progression and promote cell cycle arrest. We found that GZD824 (Olverembatinib), a broad-spectrum kinase inhibitor for ABL1, was the only compound with different effects on condensates at high and low concentrations.

While there was a weak correlation between relative condensate- and cell counts in general (Fig. S2), 40 small molecule inhibitors with effects on condensates (FDR<0.2, as described above) had no relevant negative effects on cell counts at one or both concentrations (the cut-off for normalized cell counts [z-score] was set ≥ –1). Among these compounds with clearly no negative effects on cell proliferation, we could, however, identify both, inducers (SB415286 targeting GSK3**α/**β and MAPT) and inhibitors (PF670462 and SB239063 targeting CSNK1, Nazartinib and Afatinib targeting EGFR) of condensate counts. In sum, our screening-based approach identified small molecule kinase inhibitors inducing and inhibiting Dvl condensate formation in HEK293T^DVL2_mEos^ cells in a sensitive and reproducible fashion.

### Small molecule inhibitors modulating Dvl2 condensates affect Wnt signaling

Next, we investigated whether small molecule inhibitors that modulate Dvl2 condensate formation affect Wnt signaling. To this end, we transfected a TCF4/Wnt luciferase reporter into HEK293T^DVL2_mEos^ cells (Fig. 4A). The assay was also performed after induction of the TCF4/Wnt luciferase signal by transfection of Wnt3 (Fig. S4). Cells were then treated with the small molecule library and after 48 hours, firefly luciferase and Renilla activity was measured. After normalizing for plating and other positional effects, TCF/Wnt luciferase activity was normalized to actin-Renilla signals as z-scaled log ratio.

**Fig. 4.**
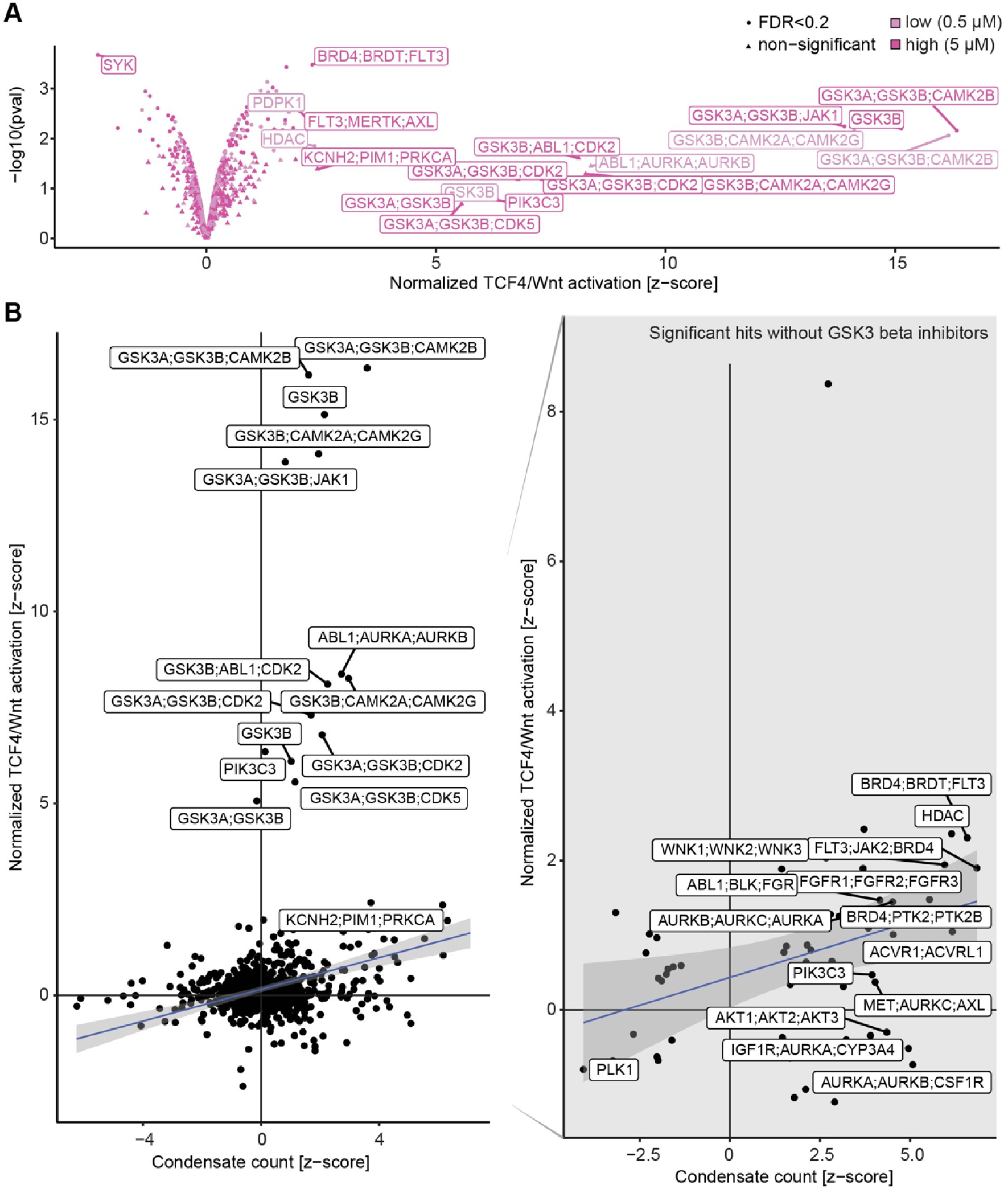
Condensate induction by small molecule kinase inhibitors correlates with Wnt signaling induction in HEK293T cells. (A) HEK293T^DVL2_mEos^ cells were seeded into white 384-well plates. After a 20 h incubation, the cells were transfected with plasmids encoding TCF4/Wnt-firefly luciferase and actin-Renilla reporters. 4 h later, the cells were treated with a custom small molecule inhibitor library at a concentration of 0.5 μM or 5 μM. Normalized TCF4/Wnt activation was determined per compound as follows: First, raw data values were normalized to the solvent (DMSO controls) per plate and replicate. Next, median row and column effects were removed from each plate in each replicate, after which the log ratio between the TCF4/Wnt-firefly luciferase reporter signals and actin-Renilla signals was computed. Finally, the ratio data was z-score scaled using the DMSO control median and the complete data variance. N=2 biological replicates for each concentration. P-values were calculated using a moderated one sample *t*.test as implemented by the lmFit function followed by an eBayes estimation. False discovery rate was estimated using multiple testing correction as defined by Benjamini and Hochberg. Selected genes are shown. The Replicate correlation was analyzed using Pearson’s correlation coefficient for high concentration (no Wnt3) = 0.69 and for low concentration (no Wnt3) = 0.82. (B) Relative condensate counts determined in the image-based screen correlate with normalized TCF4/Wnt activation determined in the TCF4/Wnt luciferase screen (PCC=0.22 without additional Wnt3 stimulation, low and high concentrations are shown). Right panel shows significant (FDR<0.2) hits while GSK3β inhibitors were excluded. Compound targets of the top 15 hits are annotated.

Without additional Wnt3 stimulus, 60 out of 643 small molecules screened modulated TCF4/Wnt luciferase activity (FDR<0.2), thereof 22 compounds induced Wnt signaling ([z-score] >1). Overall, we found a positive correlation of PCC=0.21 (p<<0.001, PCC = 0.3 when outliers are removed) of Dvl2 condensate formation and Wnt signaling activity (Fig. 4B). Top scoring hits for TCF4/Wnt luciferase induction were GSK3 inhibitors, such as CHIR-99021, AZD2858, and LY2090314, known to be strong inductors of canonical Wnt signaling (Zhong et al., 2009). In line, CK1δ/ε inhibition with strong inhibitory effects on condensate formation, showed a strong repression of Wnt3-mediated signaling induction (Fig. S4).

### The CK1δ/ε inhibitor PF670462 reduced Wnt activity and endogenous Dvl2 condensates

We then focused our further analysis on the CK1δ/ε inhibitor PF670462 (Badura et al., 2007; Cheong and Virshup, 2011), which was identified in the image-based small molecule screen to abolish endogenous Dvl2_mEos3.2 condensates (Fig. 3). PF670462 is known to bind to the ATP-binding pocket of the kinase, thereby inducing a conformational change of its gatekeeper residue Met82 (Long et al., 2012).

Time-lapse analysis in HEK293T^DVL2_mEos^ cells confirmed that treatment of cells with PF670462 led to the inhibition of novel Dvl2 condensate formation and the dissolution of nearly all existing condensates within 14 hours (Fig. 5A). At the same time, PF670462 effectively blocked Wnt3-induced and Dvl2-induced TCF4/Wnt luciferase reporter activity in HEK293T^DVL2_mEos^ cells (Fig. 5B). Importantly, PF670462 inhibited Wnt signaling also upon DVL2_mEos3.2 or Wnt3 expression in HEK293T DVL1,2,3^KO^ cells (Fig. 5B (Schubert et al., 2022)), indicating that CK1δ/ε acts at the level or downstream of Dvl2.

**Fig. 5:**
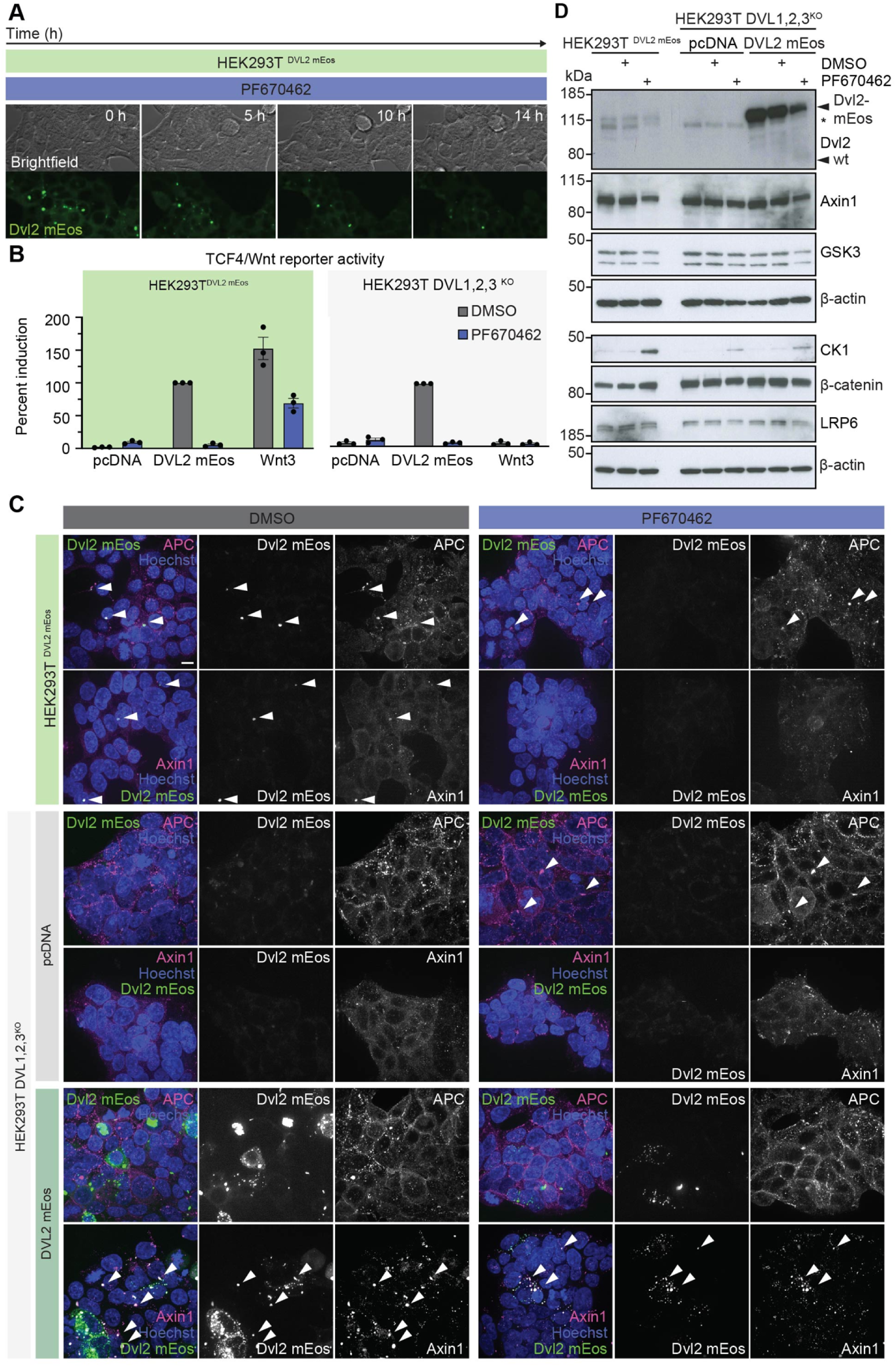
T**h**e **CK1δ/ε inhibitor PF670462 inhibits endogenous Dvl2 condensates and Wnt signaling with distinct effects on APC and Axin1.** (A) PF670462 dissolves existing Dvl2_mEos3.2 condensates and inhibits the formation of new condensates in HEK293T^DVL2_mEos^ cells. Live-cell imaging frames captured at specific time points from time-lapse videos (N=2). Cells were plated into ibidi chamber slides 24 h prior to imaging, which was performed on an A1 Nikon microscope at 15 minute intervals over 15 h under physiological conditions. The displayed images represent maximum intensity projections of four slices spanning a total of 3 µm. (B) PF670462 blocks Wnt signaling induction by Wnt3 and DVL2_mEos3.2 in HEK293T^DVL2_mEos^ cells and by DVL2_mEos3.2 expression in HEK293T DVL1,2,3^KO^ cells. Cells were transfected with pcDNA, DVL2_mEos3.2, or Wnt3 in combination with TCF4/Wnt-firefly luciferase and actin-Renilla reporter plasmids. 4 h later, the cells were treated with DMSO or 10 µM PF670462. Luciferase reporter signals were assessed 48 post-transfection. Induced reporter activity was assessed by normalization to DMSO-treated DVL2_mEos3.2 cells. Graph depicts the mean ±SEM of three separate experiments. (C) Localization analysis of Dvl2_mEos3.2, Axin1, and APC in HEK293T^DVL2_mEos^, HEK293T DVL1,2,3^KO^ pcDNA, and HEK293T DVL1,2,3^KO^ DVL2_mEos3.2 cells with and without PF670462 treatment (10 μM, 48 h). Maximum intensity projections are shown. HEK293T^DVL2_mEos^ DMSO APC (z=13) + Axin1 (z=22), HEK293T^DVL2_mEos^ PF670462 APC (z=30) + Axin1 (z=36), HEK293T DVL1,2,3^KO^ pcDNA DMSO APC (z=25) + Axin1 (z=20), HEK293T DVL1,2,3^KO^ pcDNA PF670462 APC (z=17) + Axin (z=20), HEK293T DVL1,2,3^KO^ DVL2_mEos3.2 DMSO APC (z=16) + Axin (z=21), HEK293T DVL1,2,3^KO^ DVL2_mEos3.2 PF670462 APC (z=14) + Axin1 (z=18). Exemplary images from three replicates captured in three or more points of view. Scale bar 10 μm. (D) Effects of PF670462-mediated CK1δ/ε inhibition on protein levels in HEK293T^DVL2_mEos^, HEK293T DVL1,2,3^KO^ pcDNA, and DVL2_mEos3.2 cells. 20 h after cell seeding, the specified cells were transfected with pcDNA, or DVL2_mEos. After 4 h, cells were treated with either DMSO or 10 µM PF670462. After 48 h, total cell lysates were examined *via* Western blot. Asterisk marks residual staining of Axin due to sequential use of the membrane. β-actin is used as a loading control. One of three independent experiments is shown. kDa = kilodaltons.

Interestingly, a recent publication has shown an induction of small condensates (also termed “puncta”) by PF670462 in cells overexpressing Dvl3 (Harnoš et al., 2019). Wnt ligand-induced open Dvl3 conformation showed a more uniform, i.e., “even” subcellular distribution and increased localization to Frizzled membrane receptors. CK1ε was shown to regulate Dvl3 conformation through specific phosphorylation events at the PDZ domain and its engagement with the Dvl3 C-terminus (Harnoš et al., 2019). We tested whether treatment with PF670462 in the context of an overexpressed Dvl2 similarly leads to an increase in puncta, which indeed is the case (Fig. 5C and S6). Western blotting showed differences in protein concentrations of endogenously tagged and overexpressed Dvl2_mEos3.2 protein (Fig. 5D). These results demonstrate that inhibition of CK1δ/ε leads to context-dependent effects on overexpressed cytoplasmic and endogenous centrosomal Dvl condensates, indicating that different mechanisms might be underlying the formation Dvl biomolecular condensates depending on Dvl protein concentration and nucleators present.

### The effect of CK1δ/ε inhibition on condensate composition

We next asked how CK1δ/ε inhibition affects the composition of Dvl2 biomolecular condensates. We have previously shown that large, centrosomal Dvl2 condensates showed co-localization of APC and Axin1 (Fig. 5C) (Schubert et al., 2022). Knockout of all Dvl paralogs resulted in a more uniform distribution of Axin1 and an absence of large Dvl2-positive condensates, while the distribution of APC was unaffected (Fig. 5C).

Surprisingly, we found that CK1δ/ε inhibition by PF670462 resulted in a similar phenotype in HEK293T^DVL2_mEos^ cells. When a DVL2_mEos3.2 construct was expressed in HEK293T cells with a knock-out of all Dvl paralogues (HEK293T DVL1,2,3^KO^), similarly to the analysis performed by Harnoš and colleagues (Harnoš et al., 2019), after treatment with PF670462, we observed an increase of more and smaller “puncta” per cell (Fig. 5C, for imaging at the same laser settings for comparability of signal intensities see Fig. S4). In DVL2_mEos3.2 overexpressing cells, endogenous Axin1 co-localized with Dvl2_mEos3.2 into “puncta” (Kang et al., 2022; Schwarz-Romond et al., 2007, 2005), without impacting Axin1 protein levels (Fig. 5D). Also, Axin1 protein levels were not influenced by simultaneous knockout of DVL1, 2, and 3 (Fig. 5D). In contrast, endogenous APC was not recruited to “puncta”. Interestingly, PF670462 treatment of HEK293T DVL1,2,3^KO^ cells after DVL2_mEos3.2 expression reduced the size and increased the cytoplasmic distribution of Dvl2_mEos3.2 “puncta”, while endogenous Axin1 was still found in these structures (Fig. 5C).

While we could confirm previous observations regarding the co-recruitment of Dvl2 and Axin1 applying our DVL2_mEos3.2 expression construct and endogenously tagged cell line, endogenous centrosomal Dvl2 condensates appear to be distinct structures compared to the “puncta” condensate formed upon overexpression. This difference is evidenced by overexpression-induced super-puncta possibly disassembling into smaller structures upon PF670462 treatment, which might be part of a multi-step process, whereas endogenous condensates disassemble without dispersing into structures visible by confocal microscopy.

As proposed by the Flory-Huggins theory, higher protein concentrations often promote phase separation and the assembly of condensates (Lyon et al., 2021; Flory, 1942). However, we found that Dvl2 was slightly elevated in the sorted Dvl2 condensate positive cells. To investigate whether PF670462 treatment and reduced phosphorylation by CK1δ/ε primarily induced protein degradation and thus reduced condensate formation, we treated the endogenously tagged and DVL2_mEos3.2 overexpressing cells with the CK1 inhibitor and analyzed protein levels by immunoblotting. We found that overexpressed DVL2_mEos3.2 was indeed decreased in PF670462-treated cells. At the same time, we observed only a mild effect on endogenous Dvl2_mEos3.2 (Fig. 5D). Interestingly, PF670462 treatment decreased Axin1 protein levels as observed by Western blotting, whereas DVL1,2,3^KO^ did not affect Axin1 protein levels. PF670462-treated HEK293T cells showed stabilization of CK1, as previously reported (Cheong et al., 2011; Yuan et al., 2020). CK1ε has a carboxyl-terminal extension that is rich in phosphorylation sites. Autophosphorylation of this region plays an important role in modulating kinase activity and stability, and PF670462 prevents the ability of CK1δ/ε to autophosphorylate (Cheong et al., 2011). By contrast, protein levels of β-catenin, LRP6, and GSK3β were not affected by PF670462-treatment of HEK293T cells (Fig. 5D). In summary, these findings indicate that PF670462 treatment selectively influences condensate stability and the activity of specific proteins within the Wnt signaling cascades, such as Dvl2 and Axin1, in a concentration-dependent manner, likely through the inhibition of CK1δ/ε autophosphorylation.

### A phospho-proteomic analysis of PF670462-treated cells

CK1δ/ε is known for its broad substrate specificity and plays critical roles in numerous cellular processes, including circadian rhythms and DNA repair by phosphorylating proteins such as Dvl, PER proteins, and p53 (Cheong and Virshup, 2011; Jiang, 2017). To characterize the global effects of CK1δ/ε inhibition by PF670462 on protein abundance and phosphorylation status, we performed phosphoproteomic analyses of treated and untreated HEK293T^DVL2_mEos^ cells and identified differentially abundant and phosphorylated proteins (Fig. 6A).

**Fig. 6:**
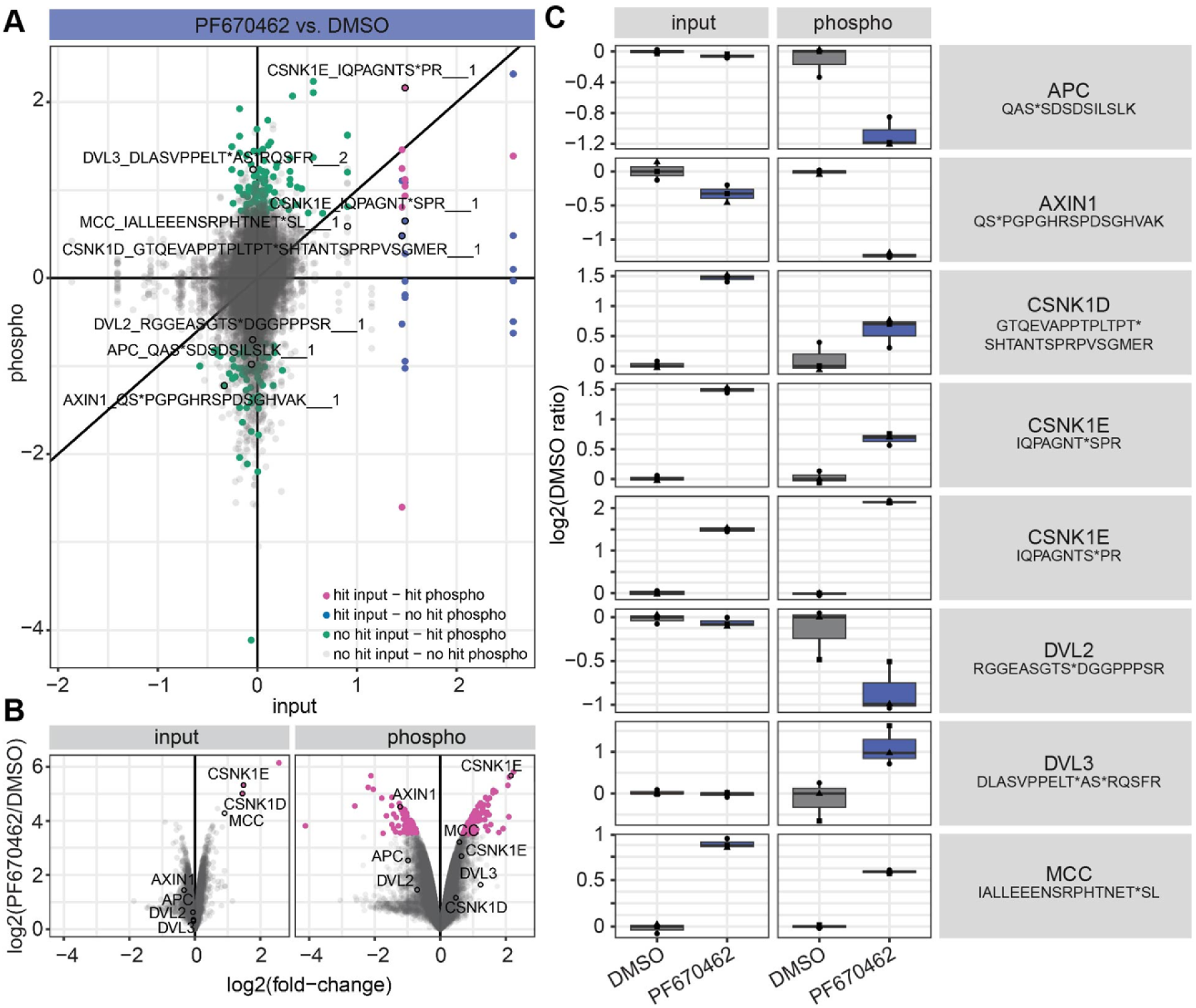
P**h**opsho**-proteomic analysis reveals differentially phosphorylated sites upon PF670462 treatment.** Fold change correlation plot (A) and volcano plot (B) of proteomic analysis reveals differential effects on protein abundance and phosphorylation status after treatment with 10 µM PF670462 for 48 h. Sample preparation and phosphopeptide enrichment followed a modified protocol based on Potel et al. (Potel et al., 2018). Details can be found in the methods section. In total, 7,102 proteins were quantified, identifying three significantly differentially expressed proteins CSNK1D, CSNK1E, and FAM199X (“hits” defined by a fold-change ≥ 50% and FDR ≤ 0.05 using limma). Additionally, 201 out of 33,175 quantified phosphopeptides were differentially regulated, including CSNK1D, CSNK1E, MCC, AXIN1. (C) Selected single plots for proteins of interest confirm effects on protein abundance observed by immunoblotting (Fig. 5D) and distinct phosphorylation status of selected phosphosites.

In total, we quantified 7102 proteins of which three proteins (CSNK1D, CSNK1E, and FAM199X) were significantly differentially expressed (“hits” defined as fold-change ≥ 50 % and FDR ≤ 0.05 with limma) and 33175 phosphopeptides of which 201 were differentially regulated (including CSNK1D, CSNK1E, MCC, AXIN1, and DDX17). Consistent with our immunoblotting results, endogenous Dvl2 protein levels were not affected by CK1δ/ε inhibition (Fig. 6B, C). However, several Dvl sites were differentially phosphorylated upon PF670462 treatment (Fig. 6C). Two less phosphorylated sites after PF670462 treatment are located in a conserved region of the Dvl paralogues.

We also observed a slight decrease in Axin1 levels, differential regulation at different phosphosites, and an increase in CK1 levels and phosphorylation (Fig. 6B, C). Furthermore, APC showed differential phosphorylation at several sites. Interestingly, another Wnt signaling-associated scaffolding protein, Mutated in colorectal cancer (MCC), was amongst the most frequently identified proteins with changes in both, total protein and phosphorylation status (Fig. 6B, C). Immunoblotting confirmed increased protein levels of MCC after treatment of HEK293T^DVL2_mEos^ cells with PF670462 (Fig. S5). MCC has been described as a negative regulator of Wnt signaling and cell proliferation (Fukuyama et al., 2008; Pangon et al., 2015), affecting cell migration and cell adhesion (Arnaud et al., 2009; Benthani et al., 2017). It has been shown to block G1 to S phase transition in NIH3T3 cells (Matsumine et al., 1996), and it is involved in DNA damage response and regulation of cell cycle arrest (Pangon et al., 2010). Recent findings indicate that MCC might be associated with the centrosome in a CK1δ/ε-dependent process (Tomaz et al., 2022). MCC and Dvl2 contain PDZ domains that mediate interactions with other PDZ domain proteins, and both also colocalize with centrosomal proteins, further supporting a potential interplay between MCC and Dvl2. Given these interactions and regulatory roles, we hypothesized that MCC might interact with Dvl2 to regulate Dvl2 condensate formation.

### CK1δ/ε inhibition by PF670462 affects cell division

We then performed GO enrichment analysis which showed an upregulation of proteins involved in the regulation of canonical and non-canonical Wnt signaling (CSNK1E, CSNK1D, MCC) (Fig. 7A). Consistent with earlier findings, we detected increased phosphorylation of proteins associated with neuron and nervous system development, including circadian rhythm regulation (Xu et al., 2005), neuronal differentiation (Bryja et al., 2007), and synaptic function (Svenningsson et al., 2002). Furthermore, we detected a downregulation of proteins associated with microtubule polymerization, depolymerization, and chromatin assembly or disassembly (Fig. 7A).

**Fig. 7:**
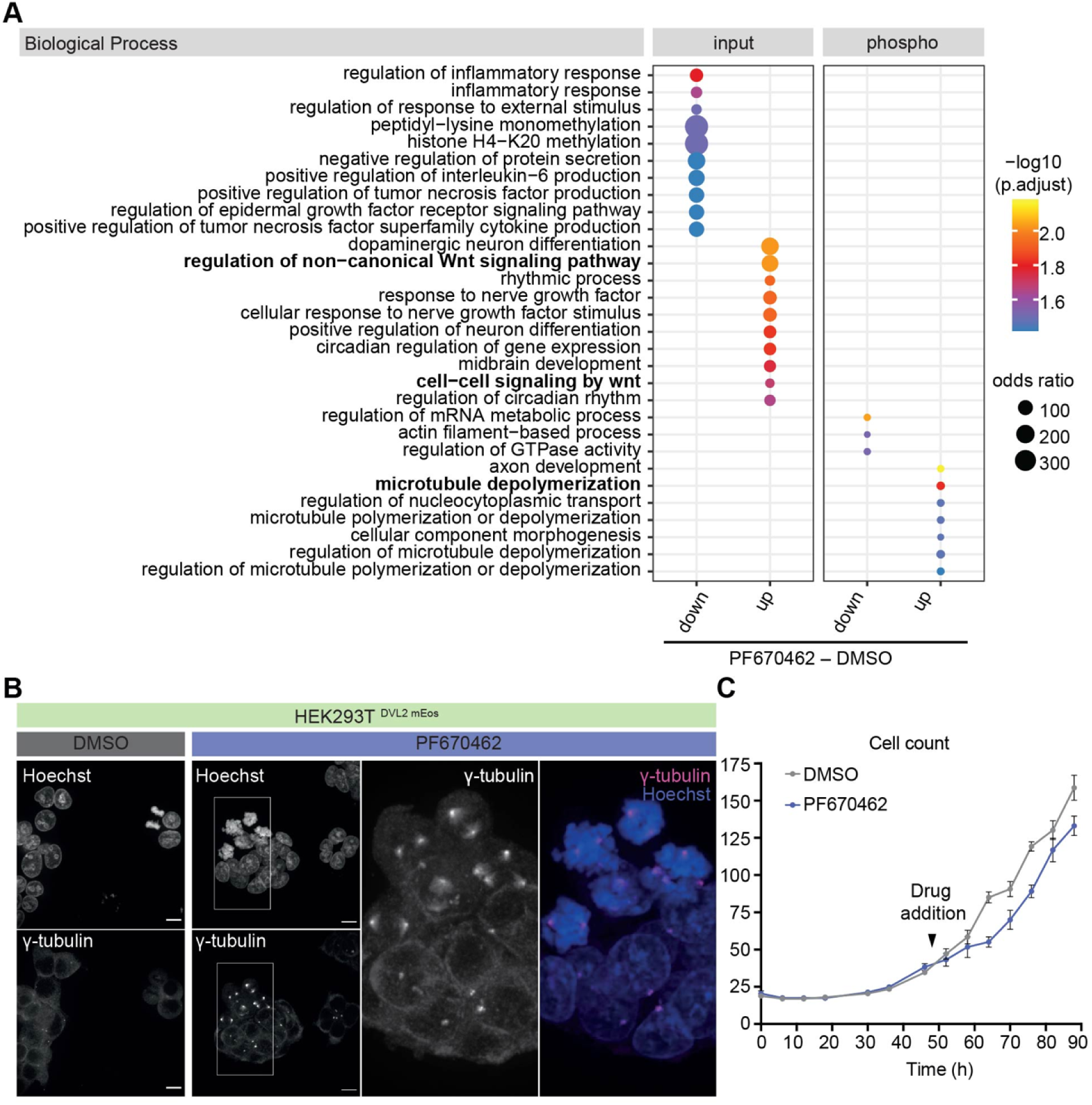
P**h**opsho**-proteomic analysis of PF670462-treated cells reveals the impact of CK1δ/ε inhibition on cell division.** (A) GO enrichment analysis demonstrates effects of CK1δ/ε inhibition on Wnt signaling and microtubule depolymerization. GO Analysis was performed as described in the methods section. The top 10 most significant terms for each category were visualized. The size of the dots represents the odds ratio and the color reflects the -log10 of the adjusted p-values. (B) Immunofluorescence staining demonstrates increased multipolar spindle formation in PF760462 vs. DMSO-treated cells. Maximum intensity projections for DMSO (z=10) and 10 µM PF670462 treatment (z=22). Exemplary images from three replicates, captured from three or more points of view. Scale bar 10 µm. (C) CK1δ/ε inhibition reduces cell proliferation. Representative Incucyte live-cell proliferation analysis. Graph shows the mean (±SEM) of measurements taken from five wells for each condition and time point. N=7.

These potential effects of CK1δ/ε inhibition on condensate formation at the centrosome (Fig. 5) and cell division (Fig. 7A) piqued our interest, as we have previously shown that depletion of all Dvl paralogs in HEK293T DVL1,2,3^KO^ cells resulted in a slight decrease of proliferation (Schubert et al., 2022). Indeed, the treatment with PF670462 induced qualitative and quantitative mitotic deficits, as observed by multipolar spindle formation (Fig. 7B) and reduced proliferation of treated HEK293T^DVL2_mEos^ cells (Fig. 7C), aligns with a role of Wnt signaling in mitosis (Augustin et al., 2017; Lin et al., 2021).

### Loss-of APC leads to an increase in Dvl2 biomolecular condensates

To further determine if proteins identified in the phosphoproteomic analysis influence the formation of Dvl2_mEos3.2 condensation, we tested whether loss-of APC, Axin2, and MCC had an impact on condensate formation. To this end, we engineered cell pools with knock-outs for AXIN1, APC, and MCC and truncated APC in HEK293T^DVL2_mEos^ cells (Fig. S6). We also generated EVI^KO^ cells as a control since they lack active Wnt signaling and show decreased condensate formation (Schubert et al., 2022) (Fig. S6). In the TCF4/Wnt luciferase reporter assay, the AXIN1^KO^, APC^KO^, and APC^trunc^ cell pools showed the expected signaling induction, while in EVI^KO^, Wnt reporter activity was abolished. MCC^KO^ did not significantly affect canonical Wnt signaling in HEK293T cells (Fig. 8A).

**Fig. 8:**
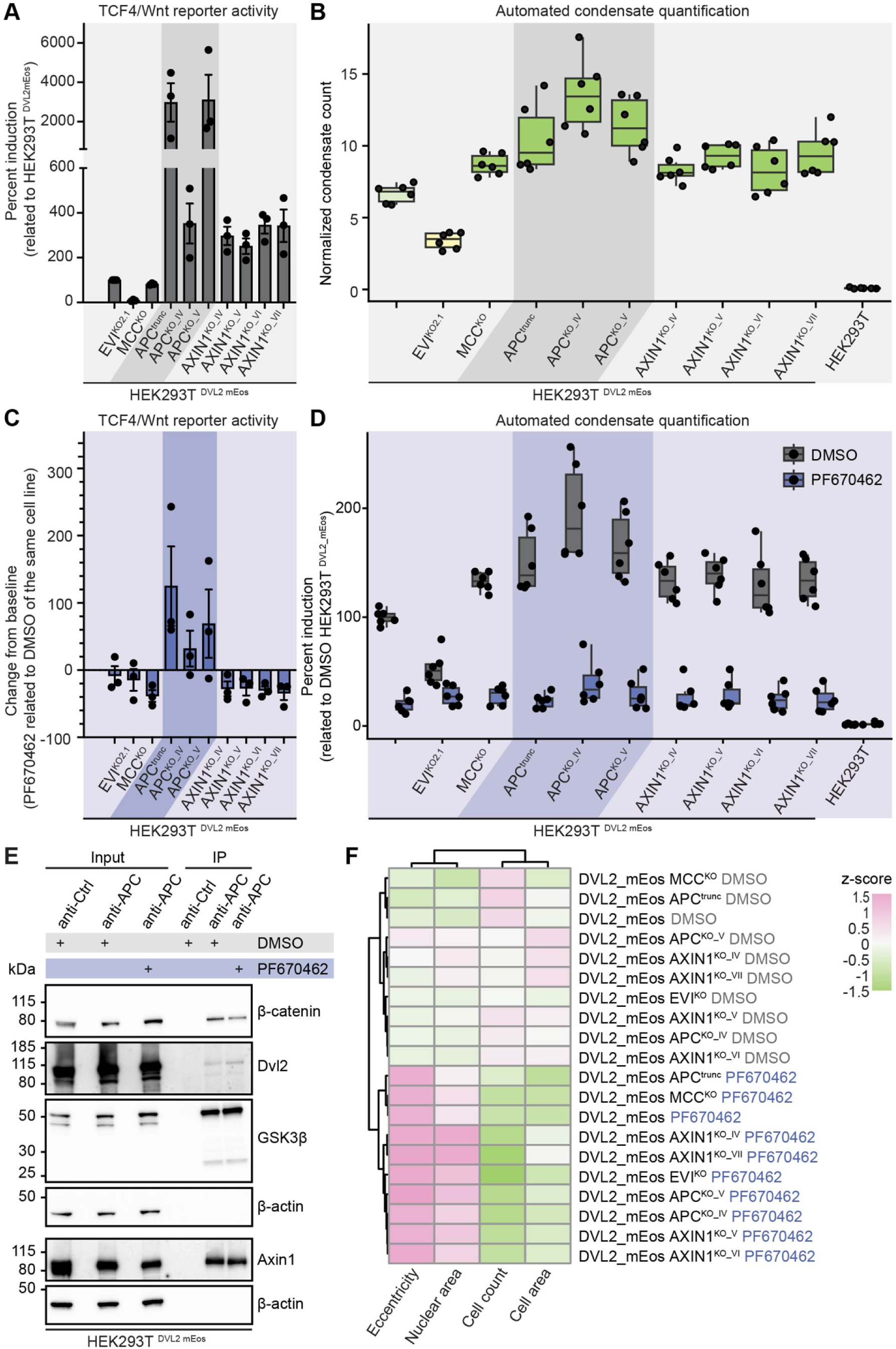
P**F**670462**-mediated CK1δ/ε inhibition induces phenotypic changes reflecting mitosis deficiency in contrast to genome engineering of Wnt scaffolds inducing Wnt signaling and condensate-induction.** (A) Validation of genome-engineered cell lines in the Wnt/TCF4 reporter assay. APC and AXIN1 modifications induce canonical Wnt signaling. Knock-out of MCC has no significant effects on Wnt signaling. The cells were transfected with TCF4/Wnt-firefly luciferase and actin-Renilla reporter plasmids. Luciferase activity was measured 48 h post-transfection. Induced reporter activity was determined by normalizing to HEK293T^DVL2_mEos^. The graph depicts the mean ±SEM of three independent experiments. Each dot represents one independent experiment. (B) Knock-out of the Wnt scaffolds APC, AXIN1, and MCC and truncation of APC induces Dvl2_mEos3.2 condensate formation. Indicated cell lines were seeded into 384-well plates, fixed after 72 h, and labeled with Hoechst and Phalloidin. Widefield images for condensate quantification were were captured using an InCell Analyzer 6000 (20x magnification). Automated image analysis was conducted as previously reported <u>(Schubert et al., 2022)</u>. The condensate count was normalized to the cell numbers in each image (condensates per 100 cells). Box plot representing the normalized condensate counts, dots indicating the average normalized condensate count per plate per 100 cells. Statistical significance was determined using the Wilcoxon signed rank test and can be found in the Table S2. (C) PF670462 treatment blocks Wnt signaling in HEK293T^DVL2_mEos^, Evi^KO^, MCC^KO^, and AXIN1^KO^ cells, but induces Wnt signaling in APC mutated cells. The cells were transfected with plasmids encoding TCF4/Wnt-firefly luciferase and actin-Renilla reporters. 4 h later, the cells were treated with DMSO, or 10 µM PF670462. Luciferase activity was measured 48 h post-transfection. Change from baseline was determined by normalizing PF670462-treatment to the DMSO control . Graph shows the mean ±SEM of three independent experiments. (D) PF670462 treatment blocks condensate formation in all cell lines. Indicated cell lines were seeded into 384-well plates and after 24 hours, the cells were treated with DMSO or 10 µM PF670462. After 48 h, the cells were fixed, stained and imaged as stated above. Box plots represent the normalized condensate counts for the indicated cell lines with dots showing the average normalized condensate count. Induction of condensates was determined by normalization to HEK293T^DVL2_mEos^ DMSO-treated cells. Statistical significance was determined using the Wilcoxon signed rank test and can be found in Table S3. (E) Immunoprecipitation of APC with and without CK1δ/ε inhibition reveals interactions independent of Dvl2 condensates. After 48 h of 10 µM PF670462 treatment, Dvl2, β-catenin, Axin1, and GSK3β still bind to APC indicating that these interactions are independent of centrosomal condensates. (F) Multiparametric feature analysis reveals clustering of PF670462-treated cells independent of Wnt scaffolds. Image analysis and feature extraction was performed as described above.

Subsequently, we quantified the Dvl2 condensates in these genetic backgrounds. Automated image-based condensate quantification revealed an increase of condensates in AXIN1^KO^, APC^KO^, APC^trunc^, and MCC^KO^ cells, with modifications of APC having the most profound effects (Fig. 8B) without having significant effects on total Dvl2_mEos3.2 protein levels (Fig. S6). Interestingly, while PF670462 treatment inhibited Wnt signaling and condensate formation in all other engineered cell lines, PF670462 in APC^KO^ and APC^trunc^ genetic backgrounds increased Wnt signaling levels as measured by the TCF4/Wnt luciferase assay while blocking condensate formation (Fig. 8C and D).

To better understand the role of APC in Dvl2 condensate formation and the differential effects of PF670462 on Wnt signaling activation in APC mutated cells, we performed an immunoprecipitation of APC in HEK293T^DVL2_mEos^ cells with and without chemical CK1δ/ε inhibition (Fig. 8E). We hypothesized that CK1δ/ε inhibition might disrupt the recruitment of Dvl2 and Axin1 to centrosomal condensates whilst their interaction with APC would still be possible. Indeed, we could still detect binding of Dvl2 and Axin1 to APC after CK1δ/ε inhibition (Fig. 8E). Imaging-based analysis of cell morphology further revealed that PF670462 treatment caused significant changes in cellular morphology, including reduced cell counts and increased nuclear size (similar to Fig. 7B, C), which were also independent of genetic background (Fig. 8F).

Taken together, our findings highlight differential functions of APC in Dvl2 condensate and Wnt signaling regulation and reveal that chemical CK1δ/ε inhibition exerts distinct effects on Wnt signaling, condensate dynamics, and cellular morphology dependent on the genetic background. While depletion of Dvl2 from centrosomal condensates correlated with cell division deficits, increased Dvl2 levels at the centrosome had no effects on cell division independent of the level of Wnt activation. Further experiments are necessary to explore the roles of Dvl2 condensates in centrosome-associated signaling and mitosis during the cell cycle.

## DISCUSSION

Biomolecular condensates have been found at several steps of the Wnt signaling pathway, including the signalosome and the destruction complex (Schaefer and Peifer, 2019; Schaefer et al., 2018, 2020; Thorvaldsen et al., 2015; Pronobis et al., 2015; Kan et al., 2020; Nong et al., 2021). Here, using image-enabled cell sorting based on phase-separated Dvl2 compartments, we analyzed condensate-positive and -negative cells. Phosphoproteomic analysis demonstrated that condensate-positive cells show an enrichment for Wnt signaling and G2/M cell cycle transition. In order to identify small molecules that modulate Dvl2 biomolecular condensate, we established an image-based high-throughput assay to quantify condensate formation. Using this assay, we performed a small molecule screen targeting kinases and focused on the CK1δ/ε inhibitor PF670462 which blocked Wnt signaling and endogenous Dvl2 condensate formation. Through genome engineering of other Wnt pathway components, we epistatically mapped the inhibitor’s effect on the formation of biomolecular condensates.

Open questions in condensate biology include understanding the mechanisms that control their formation, their molecular composition, and their functional roles in cellular processes (Lyon et al., 2021). In addition, a better understanding may also provide opportunities for targeting condensates to regulate cellular processes, leading to therapeutic interventions in diseases with aberrant condensate dynamics, such as cancer and neurodegenerative diseases (Mullard, 2019; Chakravarty et al., 2022; Lyon et al., 2021).

### Establishment of image-based screens for the analysis of endogenous Dvl2 condensates

To identify cell states and analyze perturbations in an unbiased manner, we established image-based sorting algorithms for the proteomic analysis of cells displaying endogenous Dvl2 condensates. The goal of these experiments was to characterize the states of cells with and without condensates. We next developed a high-throughput imaging assay for small molecule screens based on the decrease or increase of condensates in cells. This high-content phenotypic screen represents a novel approach to condensate biology, allowing the direct identification of perturbations with an impact on phase separation in a native cellular context. Our image-based profiling assay enables multi-parametric phenotypic analyses of kinases and pathways, providing new insights into small molecule effects on condensates and opening avenues for drug discovery, either by targeting condensates required for specific functions or as a counter-screen to eliminate small molecule hits that may have unwanted side effects such as the nonspecific pertubation of condensate formation (Mullard, 2019).

### Biomolecular condensates in Wnt signaling

Condensates in Wnt signaling pathways have attracted attention because of their potential involvement in the spatial and temporal regulation of signaling outcomes. However, what mediates and modulates condensates in Wnt signaling pathways has remained so far rather unknown. As previously shown, overexpression of Dvl induced the formation of “puncta” demonstrating the concentration dependence of LLPS (Smalley et al., 2005; Schwarz-Romond et al., 2005; Kang et al., 2022). In the sorted condensate positive cell pool, Dvl2 was slightly enriched. However, our results show that CK1δ/ε inhibition by PF670462 prevented the formation of Dvl2 condensates, whereas endogenous Dvl2 protein levels remained rather stable. Interestingly, Axin1 condensates disappeared after knockout of DVL1, 2, and 3 at stable Axin1 protein levels, whereas reduced Axin1 condensates after PF670462 treatment were associated with reduced Axin1 phosphorylation and Axin1 protein. This demonstrates that Dvl2 protein levels alone are not indicative of the protein’s ability to form condensates, but that post-translational modifications, protein-protein interactions, and the cellular context also modulate Dvl2 condensation.

Endogenous studies of the destruction complex support the interplay between Wnt signaling, cell cycle regulation, and condensates. Lach et al. demonstrated the nucleation of the destruction complex through LLPS at the centrosome (Lach et al., 2022b), reinforcing the importance of the cellular state in condensate dynamics. Wnt signaling’s pivotal role in cell division and the maintenance of genomic stability has been observed by many researchers (Stolz and Bastians, 2015; Niehrs and Acebron, 2012; Bryja et al., 2017; Habib and Acebrón, 2022; Augustin et al., 2017). However, the exact interplay of Wnt signaling and mitosis needs further investigation. Interphase centrosomal condensates have been shown to be involved in organizing microtubules and preparing for mitosis (Woodruff et al., 2017). Before mitosis, these condensates typically disappear as the cell transitions to the mitotic phase streamlining spindle formation and chromosome segregation, ensuring accurate cell division. Interestingly, we observed endogenous Dvl2 translocating to such centrosomal interphase condensates in response to Wnt activation and disappearing from these structures during mitosis (Schubert et al., 2022). In support of our data, Kikuchi and colleagues demonstrated that overexpressed Dvl2 translocates to the centrosome regulating spindle orientation in G2/M (Kikuchi et al., 2010). Also in G2/M, Cervenka and colleagues demonstrated that overexpressed Dvl2/3 can function as a scaffold for linker proteins at the centrosome (Cervenka et al., 2016).

In our study, loss of Dvl2 condensate by knockout of DVL1, 2, and 3 and inhibition of CK1δ/ε reduced cell proliferation, whereas condensate induction in the small molecule screen and by knockout of AXIN1 and truncation or knockout of APC had different effects on cell number, indicating that Wnt activity per se does not allow predicting the effect on cell division. In this context, it is noteworthy that Dvl proteins participate in “canonical” and “non-canonical” signaling cascades, including the Wnt-GSK3β-microtubule, Wnt/Ca2+, Wnt-aPKC (atypical protein kinase C), Wnt-RYK (related to tyrosine kinase), and Wnt-mTOR (mammalian target of rapamycin) pathways. Independent of Wnt signaling, Dvl proteins regulate several cellular processes, including cytoskeletal regulation, cell polarity, vesicle trafficking, and apoptosis (Sharma et al., 2018). With respect to the observations of Lach and colleagues, knockdown of β-catenin had no effect on Dvl2 condensation at the centrosome (Schubert et al., 2022). Interestingly, in the kinase inhibitor screen, increased condensates were associated with reduced cell number, larger nuclei, and increased cell area. However, PF670462 - while blocking condensate formation - affected cell proliferation and spindle formation. These results lead us to hypothesize that a delicate balance between Wnt signaling activity and the associated formation of Dvl2 condensates is essential for the cell to progress through the cell cycle without disruption. Further studies are needed to investigate whether the recruitment of Dvl2 -in response to Wnt signaling- is coincidental or essential, as well as for the re-location of β-catenin from the centrosome. Is there a specific “state” of the centrosome that is a prerequisite for Dvl2 condensation at the centrosome? And can modulation of the centrosome itself induce Dvl2 condensation at the centrosome - in the absence and/or presence of Wnt signaling and Dvl2 phosphorylation?

The distinct roles of APC and Axin1 in regulating Wnt condensates highlight their unique contributions to the destruction complex. Previous studies demonstrated the requirement of APC for Axin “puncta” assembly facilitating destruction complex formation (Mendoza-Topaz et al., 2011). CK1δ/ε phosphorylation of Dvl regulates the APC-containing destruction complex, influencing β-catenin stability and Wnt signaling outcomes (Morgenstern et al., 2017). Notably, APC co-expression led to an increase in the size of Axin “puncta” at reduced numbers, implying that APC drives incorporation of Axin into existing polymers rather than the nucleation of new ones and that APC polymerization is vital for initiating and stabilizing functional destruction complexes (Pronobis et al., 2015; Schaefer and Peifer, 2019). While the Axin-DIX domain enhances destruction complex efficiency, APC oligomerization can compensate for Axin absence by forming condensates (summarized in (Schaefer and Peifer, 2019)). Axin-Dvl heteropolymerization disrupts Axin homopolymerization and its interaction with APC, impairing complex assembly (Fiedler et al., 2011). Kang et al. recently showed at endogenous protein levels that Axin undergoes LLPS to facilitate the assembly of the destruction complex. At the same time, Dvl2 inhibits Axin’s LLPS, highlighting a critical role of Dvl2 LLPS in balancing “signalosome” and “destruction complex” formation (Kang et al., 2022). They also showed that the recruitment of Axin1 depends on the polymerization of Dvl2 at the membrane, with Dvl2’s membrane association being more stable than Axin1’s (Kang et al., 2022). Raising the Axin concentration from 100 to 1,000 nM reduced the intensity of Dvl2 “puncta” (Kang et al., 2022), in line with our observation of induced Dvl2 condensate counts in AXIN1^KO^ cells. Our results support the idea that APC and Axin, despite having overlapping functions, also possess unique roles that are critical for the destruction complex’s effectiveness, such as Axin’s interaction with GSK3 and APC’s facilitation of β-catenin transfer to the E3 ligase (Pronobis et al., 2015, 2017; Nakamura et al., 1998). In our study, Axin1 co-localized with centrosomal Dvl2 condensates and was recruited to overexpression “puncta” at stable protein levels. In DVL1,2,3^KO^ cells, Axin1 protein levels were not affected and AXIN1^KO^ did not affect Dvl2 levels. Axin1 was not observed in centrosomal condensates in DVL1,2,3^KO^ cells. In contrast to Dvl2 for Axin1 condensation, the presence of Axin1 appears unnecessary for Dvl2 condensation. APC, however, can co-localize with Dvl2 in centrosomal condensates but was not recruited to overexpression “puncta”. APC’s centrosomal localization was not affected by DVL1,2,3^KO^ or PF670462 treatment. Also, Dvl2 condensation was not dependent on APC. Assuming that Dvl2 phosphorylation is the driving force that stimulates endogenous Dvl2 to form first smaller oligomers (Ma et al., 2020; Ntourmas et al., 2024) and then larger condensates, APC might even have –direct or indirect– inhibitory effects on Dvl2 LLPS at the centrosome, since Dvl2 condensates are induced in APC^KO^ and APC^trunc^ cells.

### Inhibition of CK1δ/ε regulates condensate formation at different cellular levels

The CK1δ/ε PF670462 inhibits Dvl2 condensate formation regardless of the genetic background. However, its impact on Wnt signaling varies and depends on the genetic context. In wildtype (and possibly AXIN1^KO^) cells, PF670462 inhibits CK1δ/ε-mediated phosphorylation of critical components of the Wnt pathway, including APC, Axin1, and β-catenin, thereby suppressing Wnt signaling by interfering the destruction complex. In AXIN^KO^ cells, PF670462 still suppresses Wnt signaling, e.g., through residual interactions with APC or other components. APC acts as the main scaffold in the destruction complex, facilitating the interactions of its components, such as CK1δ/ε and Axin1. In APC^KO^ and APC^trunc^ cells, the functions of the destruction complex are disrupted. Therefore, the kinase might not be effectively recruited to the complex. Consequently, PF670462 could activate Wnt signaling by disrupting alternative phosphorylation pathways, non-canonical signaling routes, or an alternative regulatory mechanism. In this context, it is noteworthy that kinases, as co-factors and substrates, can undergo LLPS – being phase-separated or being phase-separating (López-Palacios and Andersen, 2023). CK1δ and ε are predicted to hold disordered regions (positions 301-415 and 301-4016, respectively) (UniProt Consortium 2023). Inhibited autophosphorylation and stabilization of CK1 in PF670462-treated cells might modulate its recruitment to different membraneless compartments.

### Biomolecular condensates and the cell cycle

Endogenous studies of the destruction complex support the interplay between Wnt signaling, cell cycle regulation, and condensates. Lach et al. demonstrated the nucleation of the destruction complex through LLPS at the centrosome (Lach et al., 2022b), reinforcing the importance of the cellular state in condensate dynamics. Wnt signaling’s pivotal role in cell division and the maintenance of genomic stability has been observed by many researchers (Stolz and Bastians, 2015; Niehrs and Acebron, 2012; Bryja et al., 2017; Habib and Acebrón, 2022; Augustin et al., 2017). Interphase centrosomal condensates have been shown to be involved in organizing microtubules and preparing for mitosis (Woodruff et al., 2017). Before mitosis, these condensates typically disappear as the cell transitions to the mitotic phase streamlining spindle formation and chromosome segregation, ensuring accurate cell division. Interestingly, we observed endogenous Dvl2 translocating to such centrosomal interphase condensates in response to Wnt activation and disappearing from these structures during mitosis (Schubert et al., 2022). Consistent with our data, Kikuchi and colleagues demonstrated that overexpressed Dvl2 translocates to the centrosome regulating spindle orientation in G2/M (Kikuchi et al., 2010). Also in G2/M, Cervenka and colleagues demonstrated that overexpressed Dvl2/3 can function as a scaffold for linker proteins at the centrosome (Cervenka et al., 2016). Here, Dvl2 condensate loss by knockout of DVL1, 2, and 3 and inhibition of CK1δ/ε reduced cell proliferation, whereas condensate induction in the small molecule screen and through knockout of AXIN1 and APC truncation or knockout had differential effects on cell counts indicating that Wnt activity *per se* does not allow to predict the impact on cell division. In this context, it is noteworthy, that the Dvl proteins participate in “canonical” and “non-canonical” pathways, including Wnt-GSK3β-microtubule, Wnt/Ca^2+^, Wnt-aPKC (atypical Protein Kinase C), Wnt-RYK (Related to tyrosine kinase), and Wnt-mTOR (mammalian target of rapamycin) pathways. Independent of Wnt signaling, the Dvl proteins govern several cellular processes, including cytoskeletal regulation, cell polarity, vesicle trafficking, and apoptosis (Sharma et al., 2018). Regarding the observations by Lach and colleagues, the knockdown of β-catenin had no effects on Dvl2 condensation at the centrosome (Schubert et al., 2022). Our study investigated a potential “hierarchy” by epistasis interactions of Wnt scaffolds at the centrosome. The interplay between these scaffolds ensures precise Wnt signaling regulation.

Our study underscores the potential of combining high-content screening and genome engineering to uncover novel therapeutic strategies targeting phase-separated systems. By integrating advanced imaging techniques with genetic perturbations, we demonstrate how this approach can identify key molecular regulators and pathways involved in condensate formation and -function. Specifically, our findings reveal insights into selectively targeting APC-mutant cancers, paving the way for drug screening approaches tailored to genetic and cellular contexts.

## Acknowledgment

We thank K. Löffler, M. Weidmann, A. Pöttner, A. Paneva, and L. Kirchhoff for their excellent technical support. We thank the members of the Boutros lab and the teams of the Nikon Imaging Center at Heidelberg University and the Advanced Light Microscopy Facility (ALMF) at EMBL for helpful scientific discussions. Figures were in part created with the help of BioRender. A.S. has been granted a fellowship of the DKFZ “Clinician Scientist Program”, supported by the Dieter Morszeck Foundation, and is a fellow of the DKTK School of Oncology. This work was supported by a grant of the Deutsche Forschungsgemeinschaft (DFG, German Research Foundation) SFB1324 “Mechanisms and Functions of Wnt signaling” (Project No. 331351713) to M.B. AS receives support by the Else Kröner-Fresenius-Stiftung (project 2022_EKEA.189).

## Author contributions

A.S., C.S., U.E., and M.B. designed the study. A.S., O.V., F.R., T.M., M.K., and C.S. performed the image-based high-throughput assays. A.S., B.S., M.K., and T.M. performed the TCF4/Wnt high-throughput assays. C.S. and F.H. analyzed the image-based and TCF4/Wnt high-throughput assays and integrated the data sets. A.S., F.R., and U.E. performed live-cell and confocal imaging. A.S., O.V., M.K., and N.W. validated the screening results. A.S., B.S., D.G., and D.O.-R. performed image-enabled cell sorting. A.S., B.S., J.S., and F.S. performed and analyzed mass spectroscopy. A.S., D.J., U.E., and M.B. supervised the study. A.S., C.S., F.H., D.O-R., J.S., F.S., and M.B. wrote the manuscript, D.K. reviewed and edited the original draft. All authors approved the final version of the manuscript.

## Competing interest statement

M.B. received research grants from Cellzome/GSK and Merck/Darmstadt unrelated to this study.

## MATERIAL AND METHODS

### Cell culture

HEK293T cells were sourced from ATCC (CRL-11268, Manassas, USA) and authenticated through SNP profiling (Multiplexion, Heidelberg, Germany). The cells were maintained in Dulbecco’s Modified Eagle’s Medium from Thermo Fisher Scientific (Gibco DMEM, 41965062, Waltham, USA) supplemented with 10% fetal bovine serum (FCS, F7524-500 ML, Sigma Aldrich, St. Louis, USA). Testing for mycoplasma contamination was performed on a regular basis.

### Transfections

TransIT-LT1 Transfection Reagent (731-0027, Mirus Bio, Madison, USA) was used for plasmid transfection according to the manufacturer’s instructions. Plasmids are listed below.

### Generation of CRISPR knock-out cells

The development of multitargeting sgRNAs for the DVL genes and EVI/wntless were previously reported (Voloshanenko et al., 2017; Glaeser et al., 2018; Schubert et al., 2022). SgRNAs targeting AXIN1, APC, and MCC were designed with the E-CRISP sgRNA design tool (Heigwer et al., 2014). Oligonucleotides were synthesized by Eurofins Genomics (Ebersberg, Germany), annealed, and subsequently cloned into the PX459 vector as outlined in earlier studies (Voloshanenko et al., 2017). HEK293T cells were transfected with plasmids encoding the sgRNAs and Cas9 and selected with 1-2 μg/ml puromycin (P9620, Sigma Aldrich, St. Louis, USA) for 48-72 hours. The resulting cell pools were cultured for a minimum of 5 days before proceeding with further experiments.

### Image-enabled cell sorting and time-lapse experiments

Cells were resuspended in 2% FSC 0,5 mM EDTA PBS (FACS buffer) and filtered through a 35-μm cell strainer to prevent clumping. Cell suspensions were stained with 1 μg/ml of 4′,6-diamidino-2-phenylindole (DAPI), analyzed, and sorted using the BD CellViewTM Imaging Technology (BD Life Sciences, New Jersey, USA) as previously described (Schraivogel et al., 2022). Single viable cells expressing Dvl2 condensates were identified and isolated based on the expression of mEos3.2 detected by the “Maximum Intensity 488-534/46” image parameter. Sorting was performed with a 100 μm nozzle, frequency of 34 kHz, automated stream setup by BD FACSChorus^TM^ Software, and a system pressure of 20 psi. Post-acquisition image-enabled sorting data analysis was performed using the FlowJo™ v10.8 Software (all BD Life Sciences, New Jersey, USA). Cells were sorted into 96-well clear-bottom plates (655087, Greiner, Frickenhausen, Germany) for imaging or into collection tubes for proteomics. Cells in 96-well plates were incubated under standard conditions for the indicated time points, fixed in 4% PFA, and stained with DyLight 650 Phalloidin (1:500, 12956S, Cell Signaling, Danvers, USA) and Hoechst (1:3000). For T = 0h, cells were sorted directly into PFA. Wide-field images were captured using an InCell Analyzer 6000 (GE Healthcare, Buckinghamshire, United Kingdom) with a 20x objective.

### Proteomics

Sample preparation and phosphopeptide enrichment were performed following the approach described by Potel et al. (2018) with modifications. Cell lysis was performed in a buffer containing 100 mM TRIS-HCl, 7 M urea, 1% Triton, 1 mM magnesium chloride, 5 mM Tris(2-carboxyethyl)phosphine hydrochloride, 55 mM 2-chloroacetamide (all from Sigma Aldrich, St. Louis, USA), 10 units/mL DNase I (EN0521, Thermo Fisher Scientific, Waltham, USA), as well as PhosSTOP and Complete Mini EDTA-free protease inhibitors (4906837001 and 4693159001, Merck Millipore, Burlington, USA). DNA was fragmented using sonication (10 seconds on, 60 seconds cooling on ice, repeated three times) until the sample viscosity was visibly reduced. Cellular debris was pelleted by centrifugation at 13,000 g for 10 minutes at 4°C. The resulting supernatant was treated with 1% Benzonase (1.01695.0001, Merck Millipore, Burlington, USA) and incubated at room temperature for 30 minutes. Protein (approximately 500 µg) was precipitated using the chloroform/methanol method. The precipitate was digested overnight with trypsin (Thermo Fisher Scientific, Waltham, USA) at a 1:20 enzyme-to-protein ratio in a buffer containing HEPES (pH 8.5), 1% sodium deoxycholate, 5 mM Tris(2-carboxyethyl)phosphine hydrochloride, and 30 mM 2-chloroacetamide (all from Sigma Aldrich, St. Louis, USA). Digestion was halted by adding trifluoroacetic acid (TFA) achieving a final concentration of 1%. Sodium deoxycholate was precipitated at room temperature for 15 minutes, and samples were centrifuged at 17,000 g for 10 minutes. Samples were desalted using an Oasis HLB 96-well plate (30 mg sorbent per well, 30 µm particle size; Waters Corporation, Milford, USA) with Buffer A (0.05% formic acid in LC-MS-grade water) and Buffer B (80% acetonitrile, 0.05% formic acid in LC-MS-grade water).

Peptides were resuspended in IMAC loading buffer (70% acetonitrile, 0.07% TFA), and a small aliquot from each sample was reserved for full proteome analysis. Phosphopeptide enrichment was carried out using an UltiMate 3000 RSLC LC system (Dionex Corporation, Sunnyvale, USA) equipped with a ProPac IMAC-10 column (4 x 50 mm, Thermo Fisher Scientific, Waltham, USA). For post-enrichment labeling with TMT reagents, phosphopeptides were released using 0.4% dimethylamine (Sigma-Aldrich, St. Louis, USA). Peptides were labeled with TMTsixplex (for PF670462-treated cells) or TMT10plex (for sorted cells) Isobaric Label Reagents (Thermo Fisher Scientific, Waltham, USA) following the instructions of the manufacturer. Briefly, 0.8 mg of reagent was dissolved in 42 µL acetonitrile, and 4 µL of the stock solution was added to the peptides. Samples were incubated at room temperature for 1 hour, followed by quenching with 5% hydroxylamine for 15 minutes. Subsequently, samples were combined, and cleanup was performed using an OASIS® HLB µElution Plate (Waters Corporation, Milford, USA). Fractionation of the TMT-labeled phosphoproteome (TMT6) and full proteome (TMT6 and TMT10) was conducted using high-pH reversed-phase chromatography on an Agilent 1200 Infinity HPLC system and a Gemini C18 column (3 µm, 110 Å, 100 x 1.0 mm, Phenomenex, Torrance, USA). A total of 48 fractions were collected during LC separation and later combined into 12 fractions. The pooled fractions were dried *via* vacuum centrifugation, reconstituted in 10 µL of 1% formic acid and 4% acetonitrile, and stored at -80°C until further analysis by LC-MS.

Proteomics data were acquired using an UltiMate 3000 RSLC nano LC system (Dionex Corporation, Sunnyvale, USA) coupled to an Orbitrap Fusion™ Lumos™ Tribrid™ Mass Spectrometer (Thermo Fisher Scientific, Waltham, USA). The LC system was equipped with a trapping cartridge (µ-Precolumn C18 PepMap 100, 5 µm, 300 µm i.d. x 5 mm, 100 Å) and an analytical column (nanoEase™ M/Z HSS T3, 75 µm x 250 mm, 1.8 µm, 100 Å, Waters Corporation, Milford, USA). Peptides were first concentrated on the trapping column at a flow rate of 30 µL/min with 0.05% trifluoroacetic acid for 4 minutes before being eluted onto the analytical column at a steady flow rate of 0.3 µL/min employing a binary solvent system. Solvent A consisted of 0.1% formic acid in water with 3% DMSO, while solvent B consisted of 0.1% formic acid in acetonitrile with 3% DMSO.

Different elution gradients were used based on the type of analysis performed. For the TMT6 full proteome, solvent B was increased from 2% to 8% over 2 minutes, followed by an increase to 28% over 72 minutes, then to 38% in 4 minutes, and finally to 80% in 3 minutes before being re-equilibrated to 2% in 5 minutes. For the TMT10 full proteome, solvent B was increased from 2% to 8% over 2 minutes, then to 28% over 42 minutes, to 40% in 4 minutes, and to 80% in 4 minutes, with re-equilibration to 2% in 4 minutes. For the TMT6 phosphoproteome, solvent B was increased from 2% to 8% over 2 minutes, then to 25% over 64 minutes, to 40% in 12 minutes, and finally to 80% in 4 minutes, with re-equilibration to 2% in 4 minutes. The TMT10 phosphoproteome was analyzed in a single run with solvent B increasing from 2% to 6% over 2 minutes, to 26% over 101 minutes, to 40% in 7 minutes, and to 80% in 4 minutes, followed by equilibration to 2% in 4 minutes.

Peptides were introduced to the mass spectrometer through a Pico-Tip Emitter (360 µm OD x 20 µm ID, 10 µm tip; New Objective, Woburn, USA) at a spray voltage of 2.4 kV, with the capillary temperature maintained at 275°C. Full mass scans were acquired in profile mode, with a resolution of 120,000. The mass range was set to 375–1400 m/z for the phosphoproteome and 375–1500 m/z for the full proteome, with a maximum filling time of 50 ms and an AGC target set to standard for the full proteome.

Data-dependent acquisition (DDA) was conducted for all samples. MS2 scans were acquired in profile mode with the Orbitrap resolution set to 30,000 for phosphoproteomes (15,000 for TMT6 full proteome). Fill times were set to 110 ms for phosphoproteomes, 54 ms for the TMT6 full proteome, and 94 ms for the TMT10 full proteome. The normalized AGC target was set to 200%, and a normalized collision energy of 36 was applied. The fixed first mass for MS2 scans was set to 110 m/z. Acquired proteomics data processing was carried out using Fragpipe v19.1 with MSFragger v3.7 for the proteome analysis of PF670462-treated cells and Fragpipe v20.0 with MSFragger v3.8 for the proteome analysis of sorted cells (Kong et al., 2017). The data were searched against the Uniprot proteome database *Homo sapiens* (UP000005640, ID9606, 20,594 entries, October 2022 release), with common contaminants and reversed sequences included. Fixed modifications considered were Carbamidomethyl (C) and TMT6 or 10 (K). Variable modifications included Acetyl (Protein N-term), Oxidation (M), and TMT6 or 10 (N-term), for the phosphoproteome analysis, phosphorylation on STY. A mass error tolerance of 20 ppm was applied for both MS1 and MS2 scans. Trypsin as protease with an allowance of max. two missed cleavages; seven amino acids min. peptide length; at least two unique peptides needed for identification of a protein. The FDR at the peptide and protein level was adjusted to 0.01.

### Analysis of proteomics data

The raw output files of FragPipe, including psm.tsv for phospho data and protein.tsv files for input data (Kong et al., 2017) were analyzed with R. PSMs (Peptide spectral matches) with a phosphorylation probability higher than 0.5 and proteins with at least two unique peptides were included in the analysis. Phosphorylated amino acids were annotated with an asterisk (*) placed after the phosphorylated residue in the aminoacid sequence, along with numerical labeling (1, 2, or 3) indicating the number of peptide phosphorylation sites. This sequence information was concatenated with the protein ID creating distinct phosphopeptide identifiers. The raw TMT reporter ion intensities were summed for all PSMs associated with the same phosphopeptide ID. For the total protein (input), the reporter ion intensities were taken from the protein.tsv output file. Phosphorylation signals were further normalized by input abundance. For this purpose, the reporter ion intensity for each distinct phospho ID, condition, and replicate was normalized using the formula: Norm.Intensityphospho,gene,condition = Intensityphospho,gene,condition/Intensityinput,gene,condition / median (Intensity_phospho,gene_) / median (Intensity_input,gene_) x median (Intensity_phospho,gene_). Transformed summed TMT reporter ion intensities were corrected for batch effects applying the ‘removeBatchEffects’ function from the limma package (Ritchie et al., 2015) and subsequently normalized with the variance stabilization normalization (vsn) package (Huber et al., 2002). Differential expression analysis for proteins, phosphopeptides, and normalized phosphopeptides was conducted separately using the limma package. Replicate information was incorporated as a factor in the design matrix provided to the ‘lmFit’ function. Phosphopeptides or proteins were annotated as hits if they had a FDR <5% and a fold-change of at least 100%. Candidates were identified if they showed an FDR <20% and a fold-change of at least 50%.

### Gene Ontology (GO) enrichment analysis

GO enrichment analysis was conducted using the *clusterProfiler* R package to identify enriched Cellular Component (CC), Molecular Function (MF), and Biological Process (BP) terms among the identified proteins. Protein identifiers were converted from gene symbols to Entrez IDs with the bitr() function and the *org.Hs.eg.db*database. Enrichment analyses were performed across different conditions by applying the enrichGO() function, specifying “CC”, “MF”, and “BP” as ontology categories. Odds ratios were calculated to assess overrepresentation of GO terms, comparing the gene ratios in the query list to the background using custom scripts. Significantly enriched terms (p.adjust < 0.05) were filtered and simplified using the simplify() function to reduce redundancy, with a semantic similarity threshold of 0.7. If not indicated otherwise, the top 10 significant terms per category were visualized using dot plots with ggplot2, the size of the dots representing the odds ratio and the color reflecting the -log_10_ of the adjusted p-values.

### Small molecule kinase inhibitor screen and automated condensate quantification

2,500 HEK293T^DVL2_mEos^ cells per well were seeded into DMEM 10% FCS containing 384-well black, flat-bottom µClear plates (781092, Greiner, Frickenhausen, Germany). 24 h after seeding, the cells were treated with a custom small molecule kinase inhibitor library based upon the Selleck kinase library (L1200, Selleck Chemicals GmbH, Houston, USA) based on the 2018 library version at two concentrations (high: 5 µM, low: 0.5 µM). 48 h after the treatment, cells were fixed and labeled using DyLight 650 Phalloidin (1:500) (12956S, Cell Signaling, Danvers, USA) and Hoechst (1:3000). Imaging was carried out at 20x magnification (Nikon SAC 20x objective, NA = 0.45) using an InCell Analyzer 2200 (GE Healthcare, Buckinghamshire, United Kingdom). Four fields per well were imaged. Per channel and field, a 16-bit grey-scale image (2048 × 2048 pixels) was acquired. Three individual experiments were performed as replicates. To quantify the condensates, an automated image analysis was performed with R/EBImage (Pau et al., 2010). In summary, single cells were identified based on the DNA staining (Hoechst) and F-Actin (Phalloidin). Intensity-based thresholding and region growing were applied to locate single nuclei and their corresponding cell bodies. The condensates per image were identified using intensity-based thresholding on the FITC:mEos signal inside the areas occupied by cells (Schubert et al., 2022). Morphological feature extraction involved segmenting individual nuclei, followed by generating a cell region mask via adaptive thresholding of the Phalloidin signal. Single cells were segmented by combining the nuclei mask and cell region mask through a Voronoi-based propagation algorithm (Jones et al., 2005), allowing for the calculation of numeric descriptors for cell and nuclear shape, size, staining intensity, and texture. Features were summarized per field using a trimmed mean (lower and upper 1 % quantiles were excluded) and standard deviations, resulting in a vector of 365 phenotypic features per field. Feature data was normalized per plate and log transformed using a generalized log transformation as implemented in the glog function of FitAR R package with the parameter alpha set as the 5 % quantile. All features were corrected for plate effects by the plate median and scaled . Statistically significant measurements were determined from replicate data using a moderated t-test, which tested the null hypothesis that the effect size is zero across all available measurements. The moderated t-test was performed using the lmFit and eBayes functions from the limma package in R/Bioconductor. To address multiple testing, the FDR was controlled using the Benjamini-Hochberg method, with statistically significant measurements identified at an FDR threshold of 0.2. Z-scores were averaged across replicates to compute the mean.

### Automated condensate quantification of HEK293T^Dvl2_mEos^ cells after CRISPR/Cas9-mediated knock-out of APC, AXIN1, and MCC and truncation of APC

2,500 HEK293T^DVL2_mEos^ cells and their respective CRISPR/Cas9-mediated knock-out (APC, AXIN1, and MCC) or truncation pools were seeded per well in black, clear- and flat-bottom, µClear 384-well plates (781092, Greiner, Frickenhausen, Germany). If applicable, 24 hours after cell seeding, the cells were treated with DMSO, or 10 µM PF670462 for 48 h, fixed and stained using the same protocols as described for the screening procedure. Imaging and data analysis were conducted using the same pipelines and tools as described above.

### TCF4/Wnt-reporter assay and small molecule inhibitor Wnt screen

TCF4/Wnt-luciferase assays were used to determine Wnt signaling activity in HEK293T cell lines as previously described (Voloshanenko et al., 2017). 5,000-7,500 cells per well were seeded in 384-well white, flat-bottom polystyrene plates (14239, Greiner, Frickenhausen, Germany). Transfection was carried out with 20 ng of TCF4/Wnt firefly luciferase-reporter, 10 ng of actin–Renilla luciferase-reporter, and, if applicable, 10 ng Wnt3 or DVL2_mEos plasmids 16 h after cell seeding using the TransIT-LT1 Transfection Reagent (731-0027, Mirus Bio, Madison, USA). If applicable, 24 h later, the cells were treated with small molecule inhibitors, e.g., a custom small-molecule kinase inhibitor library based upon the Selleck kinase library (L1200, Selleck Chemicals GmbH, Houston, USA) similar to the 2018 version. 48 h after the transfection, the luminescence signal was determined using the Mithras LB940 Multimode Microplate Reader (Berthold Technologies, Bad Wildbad, Germany). The TCF4/Wnt-luciferase measurements were normalized to the actin-Renilla signals. For the TCF4/Wnt-luciferase screen, two biological replicates were performed for each of the following conditions: with and without Wnt3 induction and two inhibitor concentrations (high: 5 µM, low: 0.5 µM) each.

### Analysis of TCF/Wnt Screen and data integration

Normalized TCF4/Wnt activation was determined per compound through a series of steps: First, raw data values were normalized to solvent (DMSO) controls for each plate and replicate. Next, median row and column effects were removed from each plate in each replicate. Following this, the log ratio between the TCF4/Wnt-firefly luciferase reporter signals and actin-Renilla signals was computed. Finally, the ratio data was z-score scaled using the DMSO control median and the complete data’s variance. The analysis included 2 biological replicates for each concentration. P-values were calculated using a moderated one-sample t-test as implemented by the lmFit function, followed by an eBayes p-value estimation in the R/Bioconductor package limma (Ritchie et al., 2015). The FDR was estimated using multiple testing correction as defined by Benjamini and Hochberg. Selected genes are highlighted in the figures. Replicate correlation was analyzed using Pearson’s correlation coefficient, yielding values of 0.69 for high concentration/no Wnt3, 0.82 for low concentration/no Wnt3, 0.75 for high concentration/Wnt3, and 0.72 for low concentration/Wnt3. For integration, normalized relative TCF4/Wnt-firefly was joined by compound ID with normalized relative condensate count data. Correlation was analyzed using Pearson correlation coefficient.

### Immunoblot analysis and immunoprecipitation assay

Protein was extracted with a Triton-containing lysis buffer (as described in (Voloshanenko et al., 2017)) or RIPA lysis buffer (89900, Thermo Fisher Scientific, Waltham, USA) supplemented with phosphatase and protease inhibitors (Complete Mini protease inhibitor 11836153001, Roche, Basel, Switzerland; phosphatase inhibitor 78420, Thermo Fisher Scientific, Waltham, USA) as previously reported (Schubert et al., 2022). For APC immunoprecipitations, cell extracts were incubated with either control or specific antibodies combined with Dynabeads Protein G magnetic beads (Thermo Fisher Scientific, Waltham, USA) for 14–16 hours at 4 °C. Protein amounts were determined with the Pierce BCA Protein Assay Kit (23225, Thermo Fisher Scientific, Waltham, USA). Immunoblotting was performed as previously reported <u>(Schubert et al., 2022)</u>. Signals were detected using the Amersham Hyperfilm ECL (GE28-9068-36, Cytiva, Marlborough, USA) and the COMPACT 2 NDT developer machine (PROTEC, Oberstenfeld, Germany) or the ChemiDoc MP imager (Bio-Rad Laboratories, California, USA, software version 3.0.1.14). Digital Western blot images were adjusted uniformly across the entire image for better brightness and contrast using the Image Lab software (Bio-Rad Laboratories, version 6.1.0) or Adobe Photoshop.

### Incucyte live-cell proliferation analysis

Proliferation analysis was conducted using an Incucyte live-cell imaging system as previously reported (Schubert et al., 2022).

### Live-cell time-lapse imaging

Time-lapse imaging was performed using a point-scanning A1 Nikon confocal microscope with a stage top incubation unit (TokaiHit, Fujinomiya, Japan) as previously reported (Schubert et al., 2022). In all confocal and wide-field images mEos3.2 is shown following excitation with 488 nm wavelength.

### Immunofluorescence staining

Immunofluorescence staining was performed as previously reported (Schubert et al., 2022). Images were captured using a spinning disk confocal microscope based on a Yokogawa head X1 on a Nikon Ti-E inverted microscope and the Volocity software (PerkinElmer UltraVIEW VoX). Confocal stacks were collected with a 400 nm z-step distance using a Nikon 60x water immersion objective (NA 1.27) and an EM-CCD camera (Hamamatsu C9100-02, Hamamatsu, Japan). Maximum intensity projections were created using Fiji (Schindelin et al., 2012).

### Statistical Analysis

Unless otherwise specified, statistical significance was calculated using a two-tailed Student’s t-test.

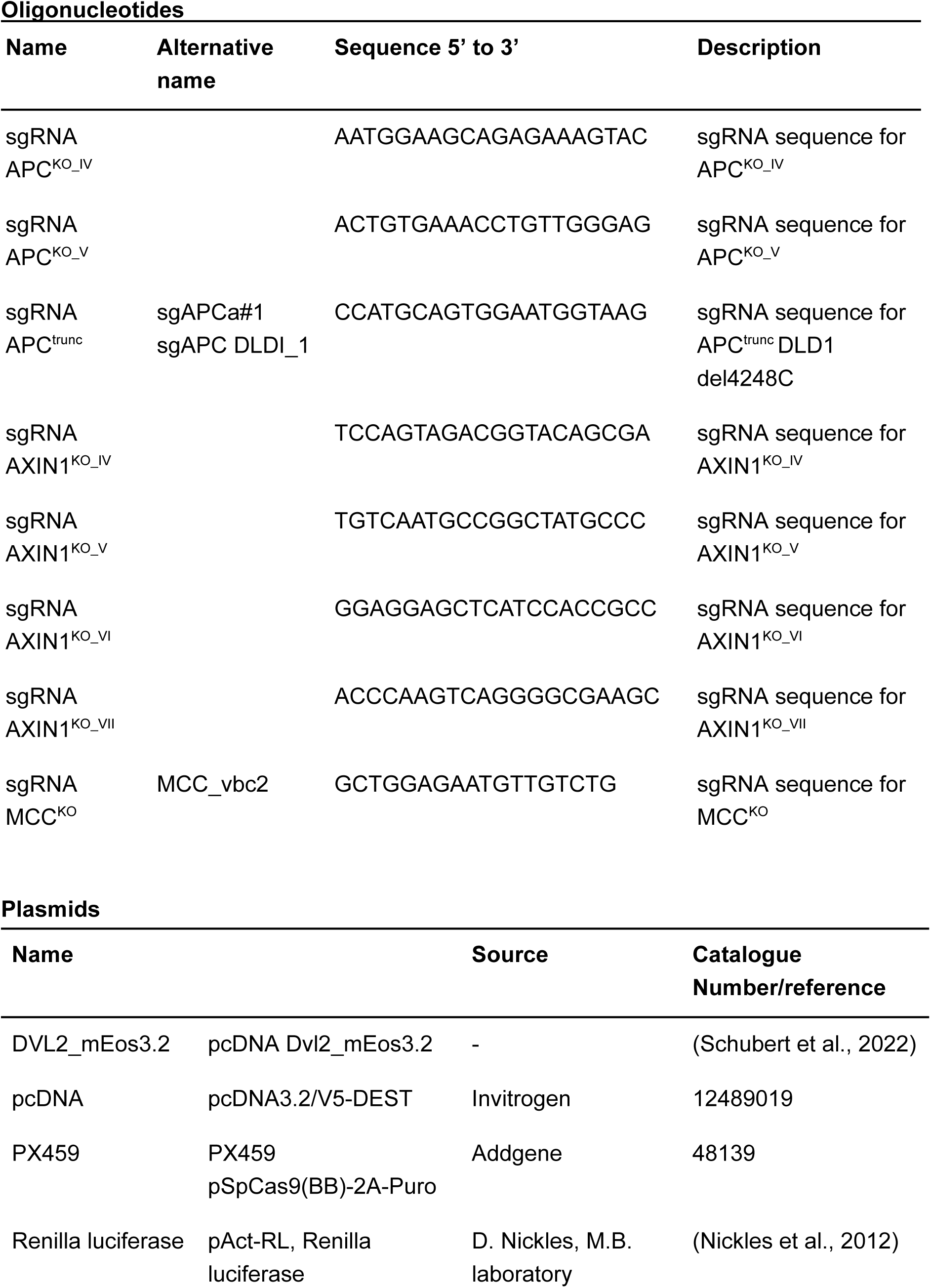
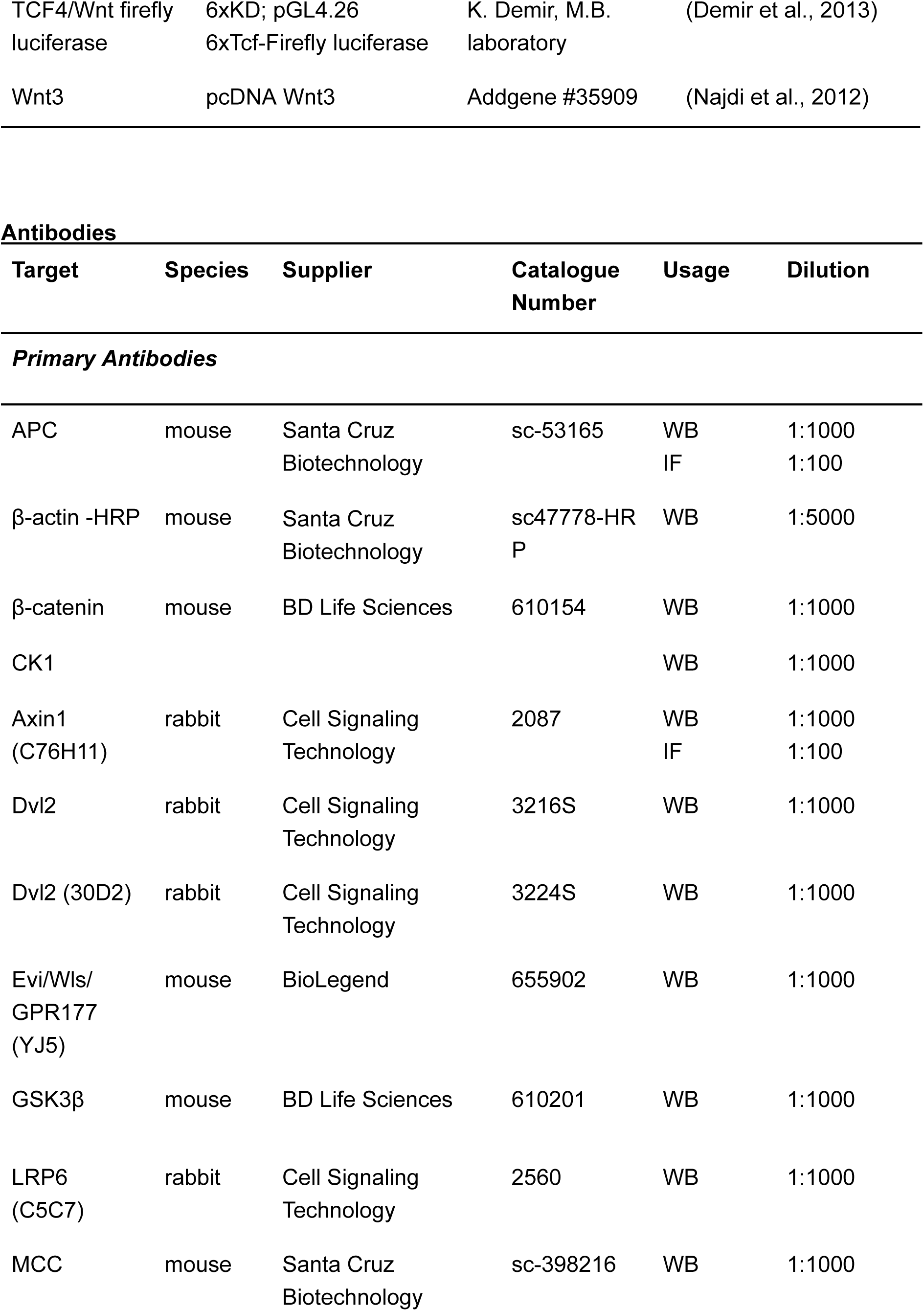
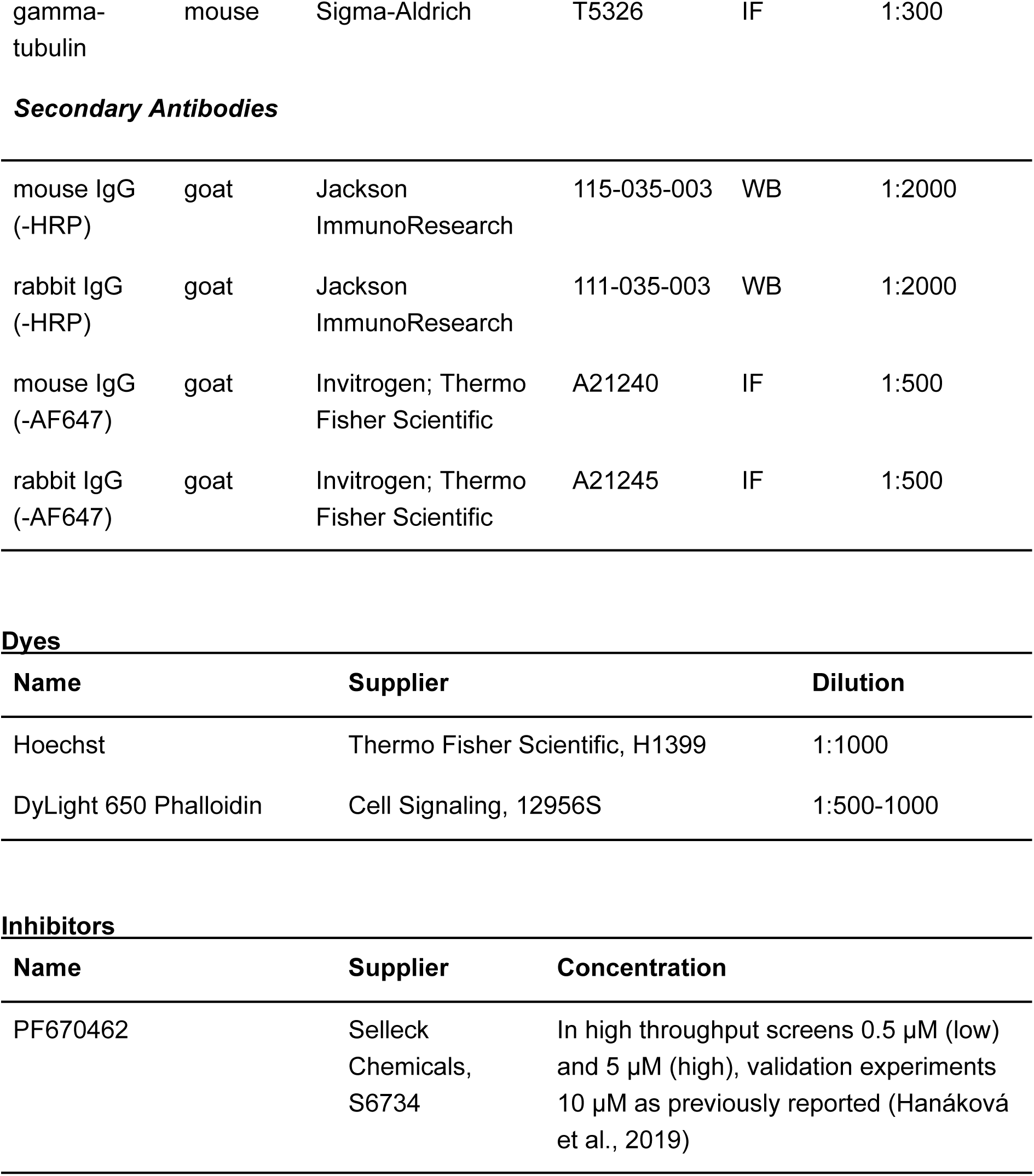

## Notes

### Competing Interest Statement

The authors have declared no competing interest.

