## Supplemental Information for "Image-based screens identify regulators of endogenous Dvl2 biomolecular condensates"

### SUPPLEMENTAL FIGURES

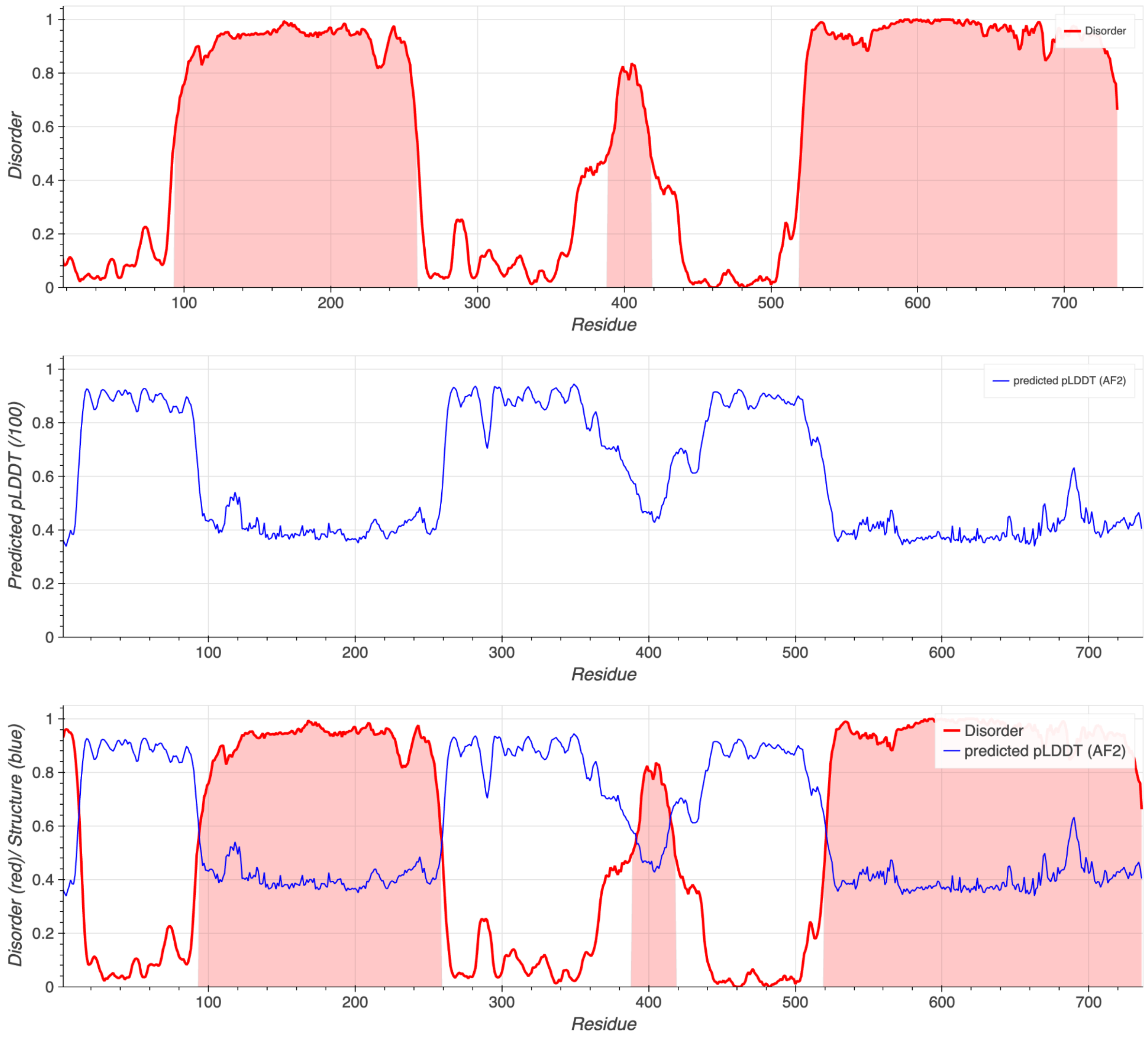


**Fig. S1: Prediction of IDRs based on metapredict.** (https://metapredict.net, accessed August 2024)(Emenecker et al., 2021, 2022).

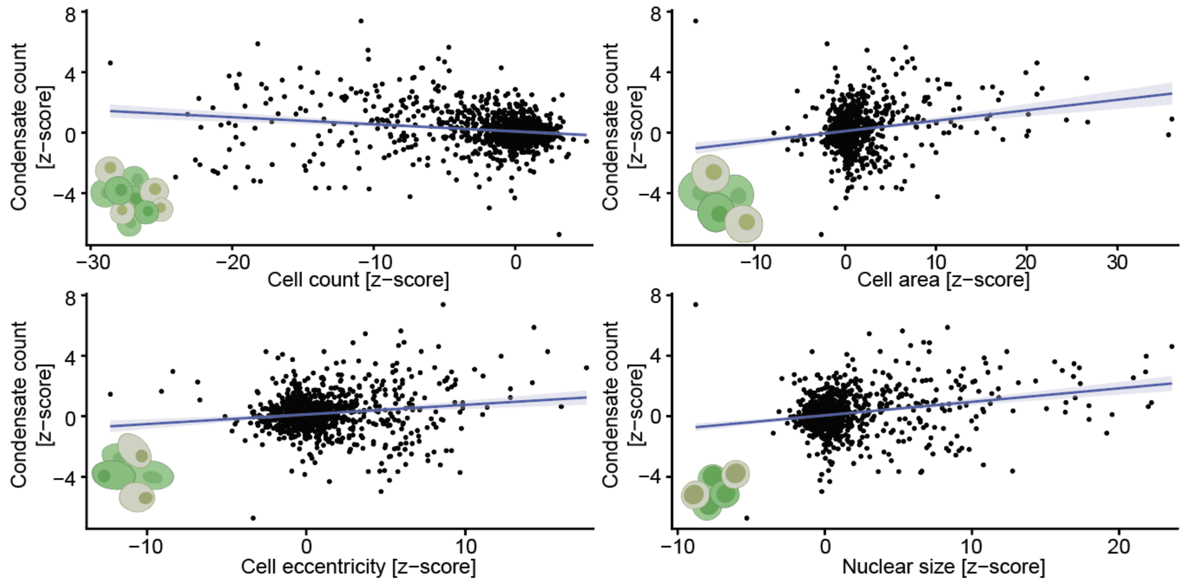


**Fig. S2: Morphological feature analysis reveals a weak correlation between condensate counts and signs of disturbed cell division.** Condensate counts correlate with cell counts, cell area, cell eccentricity, and nuclear size across compound phenotypes (PCC = -0.15, 0.19, 0.12, 0.23, respectively). To assess the correlation between condensate counts and indicated morphological features, feature pairs were analyzed using Pearson's correlation. The trend line shows a linear model fit plus the 95% Confidence interval of the fit, showing the interdependence of the variables.

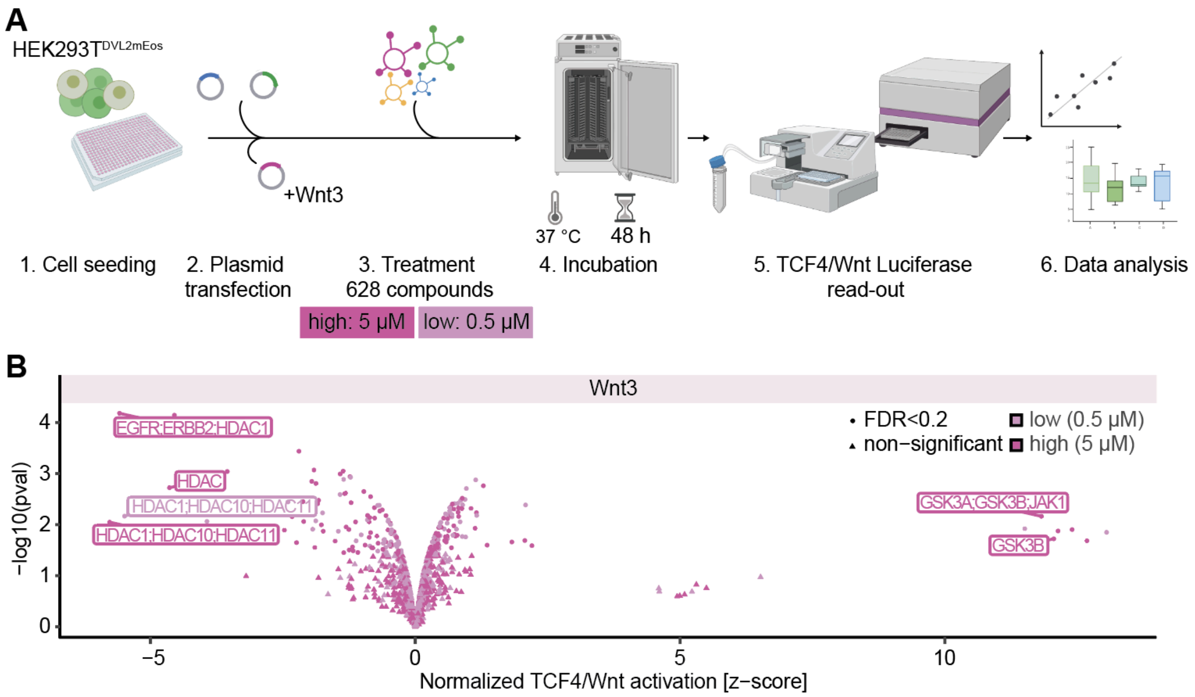


**Fig. S3: Characterization of the small molecule kinase inhibitor library for its ability to inhibit Wnt3-induced signaling.** *(A)* Schematic presentation of the TCF4/Wnt luciferase small molecule screening workflow. HEK293T$^{DVL2_mEos}$ cells were seeded into white 384-well plates. After 20 h, the cells were transfected with pcDNA Wnt3 and TCF4/Wnt-firefly luciferase and actin-Renilla-reporter plasmids. Four hours later, the cells were treated with a custom small molecule inhibitor library at a concentration of 0.5 µM and 5 µM. (B) Normalized TCF4/Wnt activation was determined per compound: first, raw data values were normalized to solvent (DMSO controls) per plate and replicate, next median row and column effects were removed from each plate in each replicate, after which the logn ratio between the TCF4/Wnt-firefly luciferase reporter signals and actin-Renilla signals was computed. Finally, the ratio data was z-score scaled using the DMSO control median and the complete data variance. N=2 biological replicates for each concentration. P-values were calculated using a moderated one sample t.test as implemented by the lmFit function followed by an eBayes estimation in the R-package limma. False discovery rate was estimated using multiple testing correction as defined by Benjamini and Hochberg. Selected genes are shown. The Replicate correlation was analyzed using Pearson's correlation coefficient for high concentration (with Wnt3) = 0.75 and for low concentration (with Wnt3) = 0.72.

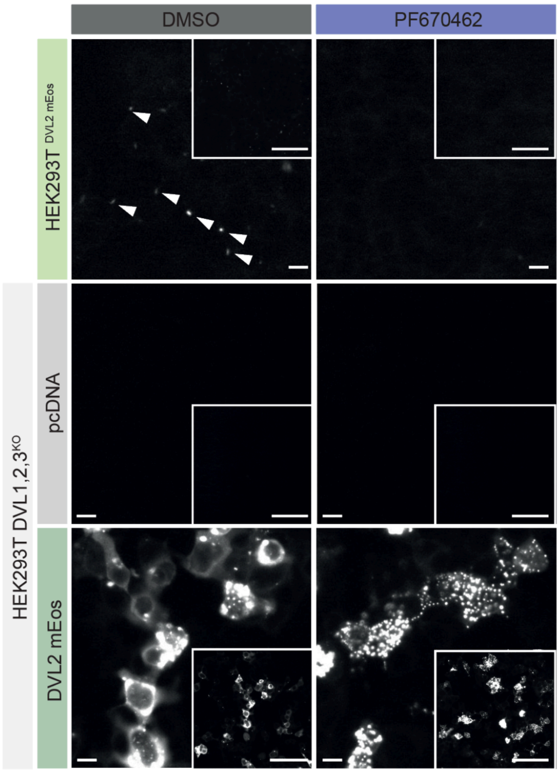


**Fig. S4: PF670462 reduces endogenous Dvl2_mEos3.2 condensates while inducing overexpression "puncta" in cells overexpressing DVL2_mEos.** Correlative wide-field images were taken of HEK293T$^{DVL2_mEos}$ and HEK293T DVL1,2,3$^{KO}$ transfected with pcDNA or pcDNA DVL2_mEos3.2 for 48 h with an InCell Analyzer 6000 (GE Healthcare, Buckinghamshire, United Kingdom). 20x magnification. For a better comparison of signal intensities between endogenous and overexpressed Dvl2, images were taken with the same laser settings for the different cell lines in contrast to the images shown in Fig. 5C. Representative images of three replicates with imaging of 3 points-of-view. Scale bar 10 µm in close-up images, 100 µm in overview images.

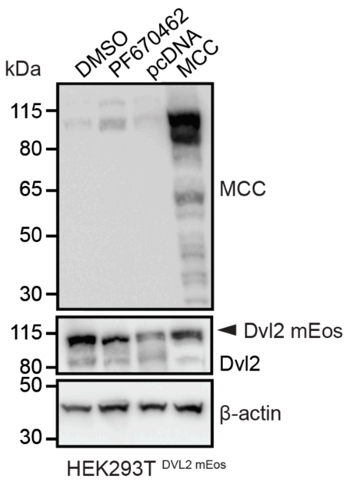


**Fig. S5: MCC protein levels are increased in HEK293T$^{DVL2_mEos}$ cells treated with PF670462.** Indicated cells were treated with either DMSO or PF670462 at a final concentration of 10 µM for 48 h. Total cell lysates were analyzed by Western blot. Cell lysate of MCC expressing cells serve as antibody control. Beta-actin serves as a loading control. One of three independent experiments is shown. kDa = kilodaltons.

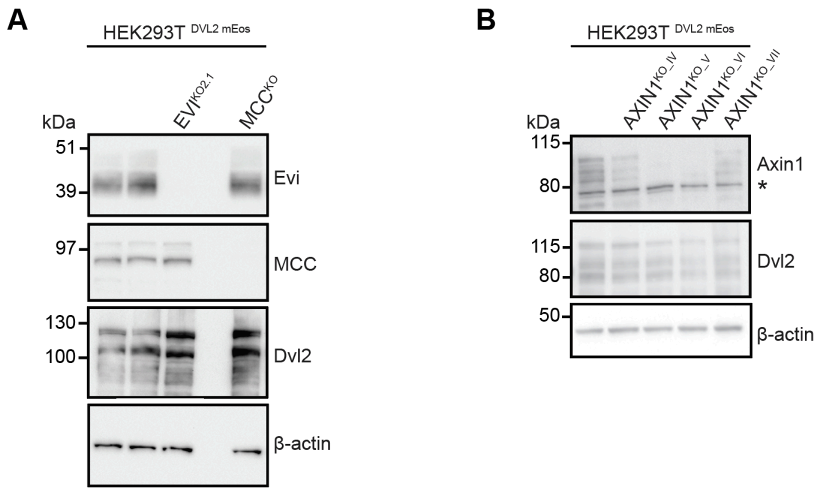


**Fig. S6: Confirmation of EVI, MCC and AXIN1 knock-out in HEK293T$^{DVL2_mEos}$ cells.** Total protein levels of Evi, MCC *(A),* and Axin *(B)* after CRISPR/Cas9-mediated knock-out of the respective genes were analyzed by Western blot. Beta-actin serves as a loading control. Asterisk marks an unspecific band. One of three independent experiments is shown. kDa = kilodaltons.

| High Concentration (5 µM) | | PCC |
|---|---|---|
| Replicate 1 | Replicate 2 | 0.43 |
| Replicate 1 | Replicate 3 | 0.38 |
| Replicate 2 | Replicate 3 | 0.81 |

| Low Concentration (0.5 µM) | | PCC |
|---|---|---|
| Replicate 1 | Replicate 2 | 0.13 |
| Replicate 1 | Replicate 3 | 0.07 |
| Replicate 2 | Replicate 3 | 0.55 |

**Table S1: Screen reproducibility was verified by evaluating the reproducibility across biological replicates.** Quality control procedures were conducted using the feature of relative condensate counts normalized to the plate median and screened replicates at both concentrations, high (5 µM) *(A)* and low (0.5 µM). Technical replicates of compounds screened in multiple wells were averaged. Pearson correlation coefficients [PCC] of the correlation between all data of respective biological replicates are shown.

| | HEK293T Dvl2_mEos | APC$^{KO_IV}$ | APC$^{KO_V}$ | APC$^{trunc}$ | Axin$^{KO_IV}$ | Axin$^{KO_V}$ | Axin$^{KO_VI}$ | Axin$^{KO_VII}$ | EVI$^{KO}$ |
|---|---|---|---|---|---|---|---|---|---|
| **APC$^{KO_IV}$** | 0.0051 | NA | NA | NA | NA | NA | NA | NA | NA |
| **APC$^{KO_V}$** | 0.0051 | 0.1981 | NA | NA | NA | NA | NA | NA | NA |
| **APC$^{trunc}$** | 0.0051 | 0.0771 | 0.4324 | NA | NA | NA | NA | NA | NA |
| **AXIN1$^{KO_IV}$** | 0.0093 | 0.0051 | 0.0093 | 0.1044 | NA | NA | NA | NA | NA |
| **AXIN1$^{KO_V}$** | 0.0051 | 0.0051 | 0.1044 | 0.3764 | 0.1044 | NA | NA | NA | NA |
| **AXIN1$^{KO_VI}$** | 0.1981 | 0.0051 | 0.1044 | 0.3089 | 1.0000 | 0.4324 | NA | NA | NA |
| **AXIN1$^{KO_VII}$** | 0.0051 | 0.0169 | 0.2450 | 0.4324 | 0.2450 | 1.0000 | 0.3764 | NA | NA |
| **EVI$^{KO}$** | 0.0051 | 0.0051 | 0.0051 | 0.0051 | 0.0051 | 0.0051 | 0.0051 | 0.0051 | NA |
| **MCC$^{KO}$** | 0.0051 | 0.0051 | 0.0169 | 0.2450 | 0.7491 | 0.3089 | 0.8562 | 0.4324 | 0.0051 |

**Table S2: Statistical analysis for Figure 8B: Knock-out of the Wnt scaffolds APC, AXIN1, and MCC and truncation of APC induces Dvl2_mEos3.2 condensate formation.** Statistical significance between the indicated knock-out pools was calculated using the Wilcoxon signed rank test and performing multiple testing correction. Shown are adjusted p-values.

| cell line | adjusted p |
|---|---|
| HEK293T$^{Dvl2_mEos}$ | 0.0026 |
| APC$^{KO_IV}$ | 0.0026 |
| APC$^{KO_V}$ | 0.0026 |
| APC$^{trunc}$ | 0.0026 |
| AXIN1$^{KO_IV}$ | 0.0026 |
| AXIN1$^{KO_V}$ | 0.0026 |
| AXIN1$^{KO_VI}$ | 0.0026 |
| AXIN1$^{KO_VII}$ | 0.0026 |
| EVI$^{KO}$ | 0.0048 |
| MCC$^{KO}$ | 0.0026 |

**Table S3: Statistical analysis for Figure 8D: PF670462 treatment blocks condensate formation in all cell lines.** Statistical significance was calculated between DMSO and PF670462 treatments of the respective cell lines using the Wilcoxon signed rank test and performing multiple testing corrections. Shown are adjusted p-values.
